# Full-spike deep mutational scanning helps predict the evolutionary success of SARS-CoV-2 clades

**DOI:** 10.1101/2023.11.13.566961

**Authors:** Bernadeta Dadonaite, Jack Brown, Teagan E McMahon, Ariana G Farrell, Daniel Asarnow, Cameron Stewart, Jenni Logue, Ben Murrell, Helen Y. Chu, David Veesler, Jesse D Bloom

## Abstract

SARS-CoV-2 variants acquire mutations in spike that promote immune evasion and impact other properties that contribute to viral fitness such as ACE2 receptor binding and cell entry. Knowledge of how mutations affect these spike phenotypes can provide insight into the current and potential future evolution of the virus. Here we use pseudovirus deep mutational scanning to measure how >9,000 mutations across the full XBB.1.5 and BA.2 spikes affect ACE2 binding, cell entry, or escape from human sera. We find that mutations outside the receptor-binding domain (RBD) have meaningfully impacted ACE2 binding during SARS-CoV-2 evolution. We also measure how mutations to the XBB.1.5 spike affect neutralization by serum from individuals who recently had SARS-CoV-2 infections. The strongest serum escape mutations are in the RBD at sites 357, 420, 440, 456, and 473—however, the antigenic impacts of these mutations vary across individuals. We also identify strong escape mutations outside the RBD; however many of them decrease ACE2 binding, suggesting they act by modulating RBD conformation. Notably, the growth rates of human SARS-CoV-2 clades can be explained in substantial part by the measured effects of mutations on spike phenotypes, suggesting our data could enable better prediction of viral evolution.

## Introduction

Over the last four years of SARS-CoV-2 evolution, the virus has accumulated mutations throughout its genome. The most rapid evolution has occurred in the viral spike: for instance, the currently dominant XBB-descended variants have 45–48 spike protein mutations relative to the earliest known strains from Wuhan in late 2019. The reason for this rapid evolution is that spike mutations can strongly affect both the virus’s inherent transmissibility and ability to escape pre-existing immunity^1–3^. A crucial aspect of interpreting and forecasting SARS-CoV-2 evolution is therefore understanding the impact of current and potential future mutations to spike.

Here we measure how thousands of mutations to the spike glycoprotein of recent SARS-CoV-2 strains impact three molecular phenotypes critical to viral evolution: cell entry, ACE2 binding, and neutralization by human polyclonal serum (**Fig. 1a**). To do this, we extend a recently described pseudotyped lentivirus deep mutational scanning system^4^ that enables safe experimental characterization of mutations throughout the spike^5,6^. We demonstrate that mutations outside the RBD can substantially impact spike binding to ACE2. We also define the mutations that escape neutralization by sera from humans who have been multiply vaccinated and also recently infected by XBB or one its descendant lineages (XBB*), and show there is appreciable heterogeneity in the antigenic impact of mutations across individuals. Finally, we show that the spike phenotypes we measure explain much of the changes in viral growth rate among different XBB-descended viral clades that have emerged in humans over the last year.

**Fig. 1:**
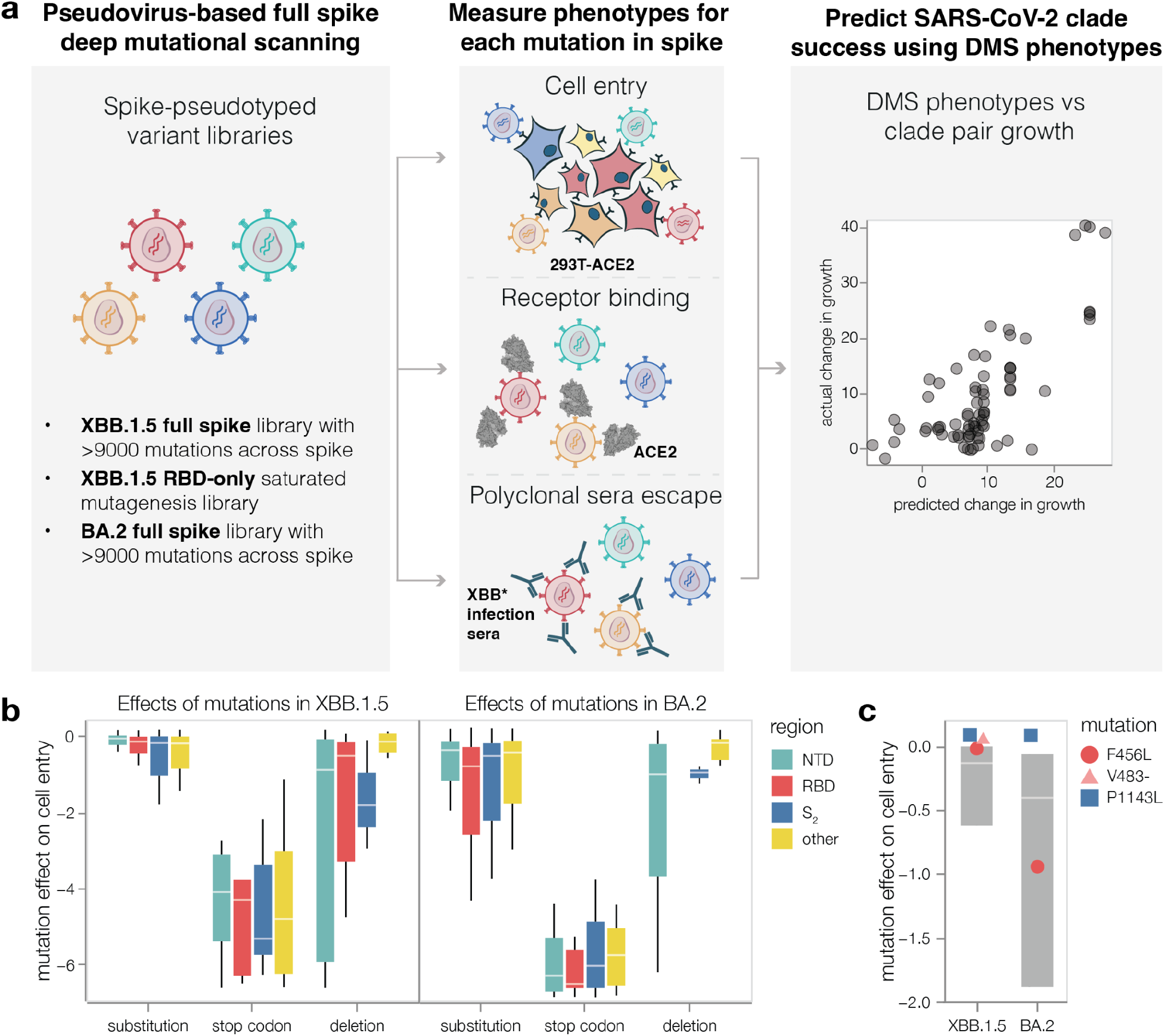
Deep mutational scanning to measure phenotypes of the XBB.1.5 and BA.2 spikes. **a,** We measure the effects of mutations in spike on cell entry, receptor binding and serum escape. We then use these measurements to predict the evolutionary success of human SARS-CoV-2 clades. **b,** Distribution of effects of mutations in XBB.1.5 and BA.2 spikes on entry into 293T-ACE2 cells for all mutations in the deep mutational scanning libraries, stratified by the type of mutation and the domain in spike. Negative values indicate worse cell entry than the unmutated parental spike. Note that the library design favored introduction of substitutions and deletions that are well tolerated by spike, explaining why many mutations of these types have neutral to only modestly deleterious impacts on cell entry. **c,** Cell entry effects of mutations F456L, P1143L and deletion of V483 relative to the distribution of effects of all substitution and deletion mutations in the libraries. Interactive heatmaps with effects of individual mutations across the whole spike on cell entry are at https://dms-vep.github.io/SARS-CoV-2_XBB.1.5_spike_DMS/htmls/293T_high_ACE2_entry_func_effects.html and https://dms-vep.github.io/SARS-CoV-2_Omicron_BA.2_spike_ACE2_affinity/htmls/293T_high_ACE2_entry_func_effects.html. The boxes in panels b and c span the interquartile range, with the horizontal white line indicating the median. For panel c, the effect of deleting V483 was not measured in the BA.2 spike.

## Results

### Design of deep mutational scanning libraries

We created mutant libraries of the spikes from the XBB.1.5 and BA.2 strains. We chose these strains because nearly all currently circulating human SARS-CoV-2 descends from either BA.2 or XBB.1.5’s parent lineage XBB^7^, and because XBB.1.5 is the sole component of the COVID-19 booster vaccine currently recommended by the WHO^8^. We wanted the libraries to contain all evolutionary accessible amino-acid mutations tolerable for spike function. We therefore included all mutations observed at an appreciable number of times among the millions of SARS-CoV-2 sequences in GISAID. In addition, we included all possible mutations at sites that change often during SARS-CoV-2 evolution or are antigenically important^3,9^, and deletions at key NTD and RBD sites. These criteria led us to target ∼7,000 amino-acid mutations in each of the XBB.1.5 and BA.2 libraries (**Extended Data Fig. 1a**). We created two independent libraries for each spike so we could perform all deep mutational scanning in full biological duplicate. The actual libraries contained between 69,000 and 102,000 barcoded spike variants with an average of 2 mutations per variant, and successfully covered 99% of the targeted mutations as well as some additional mutations (**Extended Data Fig. 1a**). To retrospectively validate that this library design strategy covered most evolutionarily important mutations, we confirmed that our libraries provided adequate coverage for high-confidence experimental measurements of nearly all mutations present in XBB or BA.2 derived Pango lineages as of the time of writing this manuscript (**Extended Data Fig. 1b**). Because the RBD is an especially important determinant of ACE2 binding and serum antibody escape^2,10^, we also made duplicate XBB.1.5 libraries that saturated all amino-acid mutations in the RBD only (**Extended Data Fig. 1a**).

### Effects of spike mutations on cell entry

We measured the effects of all library mutations on spike-mediated cell entry in 293T-ACE2 cells^4^ (**Extended Data Fig. 1c-d** and interactive heatmaps at https://dms-vep.github.io/SARS-CoV-2_XBB.1.5_spike_DMS/htmls/293T_high_ACE2_entry_func_effects.html and https://dms-vep.github.io/SARS-CoV-2_Omicron_BA.2_spike_ACE2_binding/htmls/293T_high_ACE2_entry_func_effects.html). These measurements were highly correlated between the replicate libraries for each spike, indicating the experiments have good precision (**Extended Data Fig. 1e**). The effects of mutations were also well correlated between the XBB.1.5 and BA.2 spikes (**Extended Data Fig. 1f**), consistent with prior reports that most but not all mutations have similar impacts on the spikes of different SARS-CoV-2 variants^11,12^. As expected, stop codons were highly deleterious for cell entry (**Fig. 1b**). Because our library design strategy favors functionally tolerated mutations in spike, most amino-acid mutations in our libraries just slightly impaired cell entry, and some but not all single-residue deletions were also well tolerated (**Fig. 1b**). Overall, the effects of mutations on cell entry were fairly well correlated with the effects of amino-acid mutations on viral fitness estimated from millions of natural human SARS-CoV-2 sequences (**Extended Data Fig. 1g**)^13^.

No individual mutation in either the XBB.1.5 or BA.2 spikes dramatically increased pseudovirus cell entry, though some mutations did marginally improve entry (**Fig. 1b** and interactive heatmaps linked in figure legend). One mutation that slightly improves pseudovirus entry in both XBB.1.5 and BA.2 is P1143L (**Fig. 1c**), which is found in the recently emerged BA.2.86 lineage^14^. We previously reported that mutations to P1143 also improve cell entry for BA.1 and Delta pseudoviruses^4^. The deletion mutations in our libraries are usually more deleterious for cell entry than substitutions (**Fig. 1b**); however, deletion of V483 in the RBD is well tolerated for cell entry, consistent with emergence of this mutation in the BA.2.86 variant^14^. The F456L mutation, which has emerged repeatedly in XBB clades after being rare in earlier BA.2-derived clades, is well tolerated for cell entry in XBB.1.5 but substantially deleterious in BA.2 (**Fig. 1c**).

### Both RBD and non-RBD mutations affect spike binding to ACE2

To measure how mutations in spike affect receptor binding, we leveraged the fact that the soluble ACE2 ectodomain neutralizes spike-mediated infection with a potency proportional to the strength of spike binding to ACE2^3,15^. Specifically, soluble ACE2 more potently blocks entry by spikes with mutations that increase binding to ACE2. To validate this fact, we made pseudoviruses with six different spike variants and quantified their neutralization by monomeric ACE2 (**Fig. 2a**). Compared to the BA.2 spike, the Wuhan-Hu-1 + D614G spike is neutralized less potently by soluble ACE2 consistent with its weaker ACE2 binding^16,17^, whereas four mutants of BA.2 known to have higher ACE2 binding^18^ (N417K, N417F, R493Q, and Y453F) were all neutralized more potently by soluble ACE2 (**Fig. 2a**). The quantitative neutralization by soluble ACE2 was highly correlated with previously measured monomeric RBD ACE2 affinities^17–19^ (**Fig. 2b**).

**Fig. 2:**
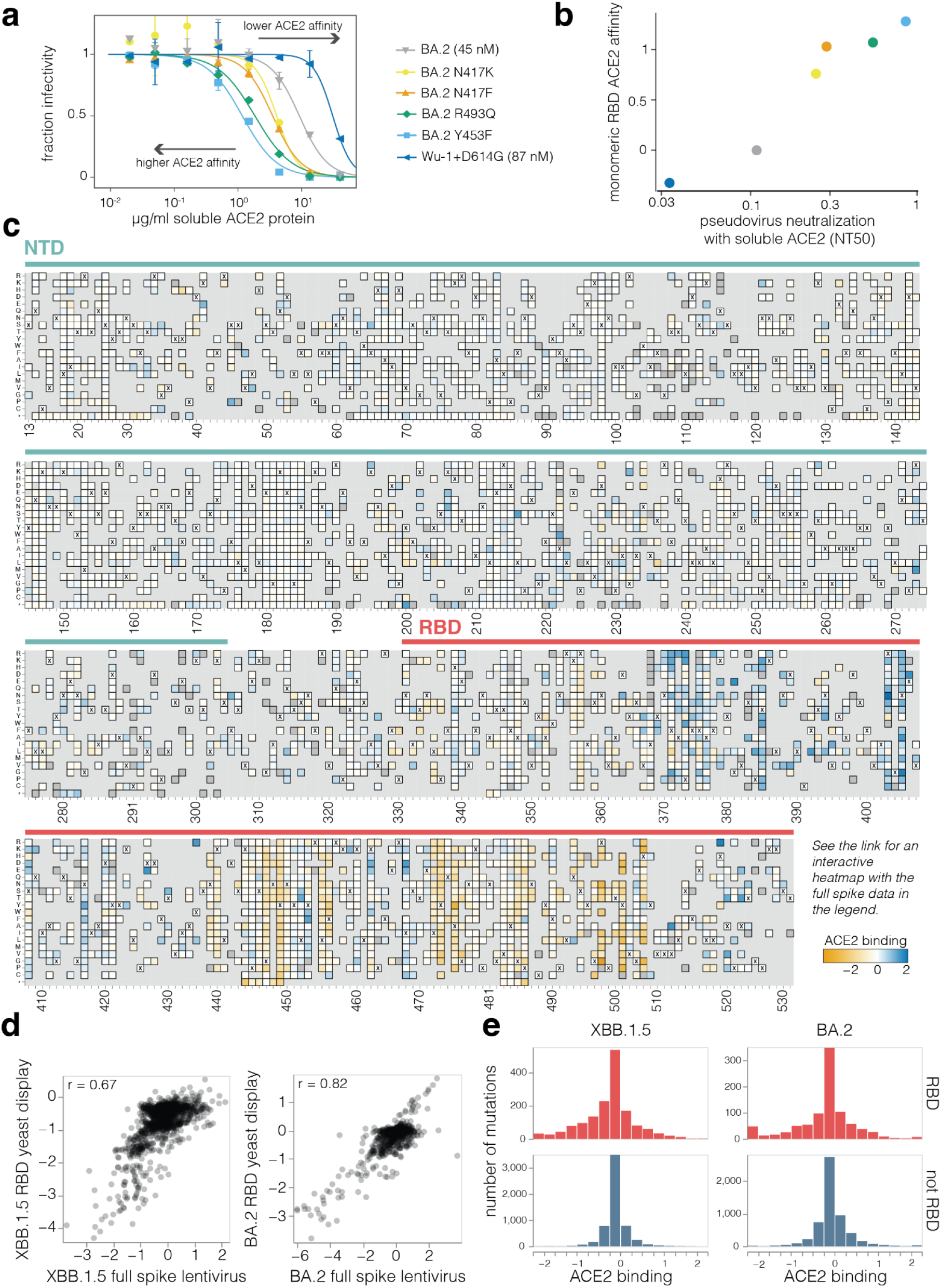
Effects of mutations on full-spike ACE2 binding measured using pseudovirus deep mutational scanning. **a,** Neutralization of pseudoviruses with the indicated spikes by soluble monomeric ACE2. Viruses with spikes that have stronger binding toACE2 are neutralized more efficiently by soluble ACE2 (lower NT50), whereas viruses with spikes with worse binding are neutralized more weakly. ACE2 affinity values measured by surface plasmon resonance for BA.2 and Wu-1+D614G are shown in brackets^17^. **b,** Correlation between neutralization NT50 by soluble ACE2 versus the RBD affinity for ACE2 as measured by titrations using yeast-displayed RBD^18^. **c,** Effects of NTD and RBD mutations on full-spike ACE2 binding as measured using pseudovirus deep mutational scanning. Mutations that enhance ACE2 binding are shaded blue, mutations that decrease affinity are shaded orange, mutations that are too deleterious for cell entry to be measured in the binding assay are dark gray, and light gray indicates mutations not present in our libraries. Interactive heatmaps showing mutational effects on ACE2 binding for the full XBB.1.5 and BA.2 spikes are at https://dms-vep.github.io/SARS-CoV-2_XBB.1.5_spike_DMS/htmls/monomeric_ACE2_mut_effect.html and https://dms-vep.github.io/SARS-CoV-2_Omicron_BA.2_spike_ACE2_affinity/htmls/monomeric_ACE2_mut_effect.html. Note that a few sites are missing in the static heatmap in this figure due to lack of coverage or deletions in the XBB.1.5 spike; see the interactive heatmaps for per-site numbering. **d,** Correlations between the effects of RBD mutations on ACE2 binding measured using the pseudovirus-based approach (this study) and yeast-based RBD display^18,20^. **e,** Distribution of effects of individual mutations on full-spike ACE2 binding for all functionally tolerated mutations in our libraries, stratified by RBD versus non-RBD mutations. Note that effects of magnitude greater than two are clamped to the limits of the plots’ x-axes.

Using this approach, we measured how mutations across both the XBB.1.5 and BA.2 spikes affect ACE2 binding (**Fig. 2c** and interactive plots at https://dms-vep.github.io/SARS-CoV-2_XBB.1.5_spike_DMS/htmls/monomeric_ACE2_mut_effect.html and https://dms-vep.github.io/SARS-CoV-2_Omicron_BA.2_spike_ACE2_binding/htmls/monomeric_ACE2_mut_effect.html). Because our assay is based on ACE2 neutralization there are several distinct mechanisms that could affect what we refer to as ACE2 binding: direct changes in 1:1 RBD-ACE2 binding affinity^18,20^, changes in spike that modulate the conformation of the RBDs, such as up/down movements^21,22^, and ACE2-induced shedding of the S_1_ subunit^23,24^.

The effects of RBD mutations on ACE2 binding to the spike measured using pseudovirus deep mutational scanning correlate well with previously reported measurements from RBD yeast-display for both XBB.1.5 and BA.2 (**Fig. 2d**)^20^. We also measured ACE2 binding for the XBB.1.5 pseudovirus libraries with saturating RBD mutations using both monomeric and dimeric soluble ACE2. The RBD-only pseudovirus measurements were highly correlated with the full-spike library measurements (**Extended Data Fig. 2a**), and the measured values were highly similar for monomeric versus dimeric soluble ACE2 (**Extended Data Fig. 2b**). Importantly, ACE2 binding and pseudovirus cell entry are distinct properties, with no strong correlation between these properties among tolerated mutations (**Extended Data Fig. 2c**) – likely reflecting the fact that cell entry can be limited by factors unrelated to receptor binding, especially in target cells expressing moderate to high levels of ACE2 like those used in our experiments.

A striking observation from the deep mutational scanning is that some mutations outside the RBD appreciably affect binding to ACE2 (**Fig. 2c,e** and **Extended Data Fig. 2d**). To validate these findings, we used mass photometry to measure binding of the soluble native ACE2 dimer to the spike ectodomain trimer (**Fig. 3a**). Mass photometry measures protein-protein interactions in solution by detecting changes in light scattering that are proportional to protein molecular mass^25^, which allows us to detect binding of one or more ACE2 molecules to the spike (**Fig. 3a**). We produced prefusion-stabilized HexaPro BA.2 wildtype spike, along with mutants that our deep mutational scanning experiments showed to increase ACE2 binding, and performed mass photometry in the presence of a series of ACE2 concentrations (**Fig. 3a-b, Extended Data Figs. 3, 4 and 5**). As expected, we observed better and worse ACE2 binding for RBD mutations that have been previously identified to either increase (R493Q) or decrease (R498V) ACE2 engagement, respectively^18^ (**Fig. 3b**, left panel). Furthermore, we detected increased ACE2 binding to BA.2 spike trimers harboring S_1_ subunit mutations (in NTD, RBD, and SD1 domains) that our deep mutational scanning indicated had better binding (**Fig. 3b** middle panel, **Extended Data Fig. 3**). For these S_1_ mutants, the increase in binding was particularly pronounced for the second ACE2 molecule (**Extended Data Fig. 5**). However, two mutations to the BA.2 S_2_ subunit found to increase binding to ACE2 in our deep mutational scanning (A701M and D950N) did not lead to increased ACE2 binding detectable by mass photometry (**Fig. 3b** right panel, **Extended Data Figs. 3 and 5b-c**). Notably, mutations at both of these S_2_ sites were previously reported to affect spike fusion^26,27^. Unlike the spikes in deep mutational scanning experiments, spikes used in mass photometry experiments are pre-fusion stabilized by mutations in fusion machinery^28^. These modifications to spike may limit the propagation of long-range allosteric changes induced by S_2_ subunit mutations, possibly explaining the discrepancy between deep mutational scanning and mass photometry.

**Fig. 3:**
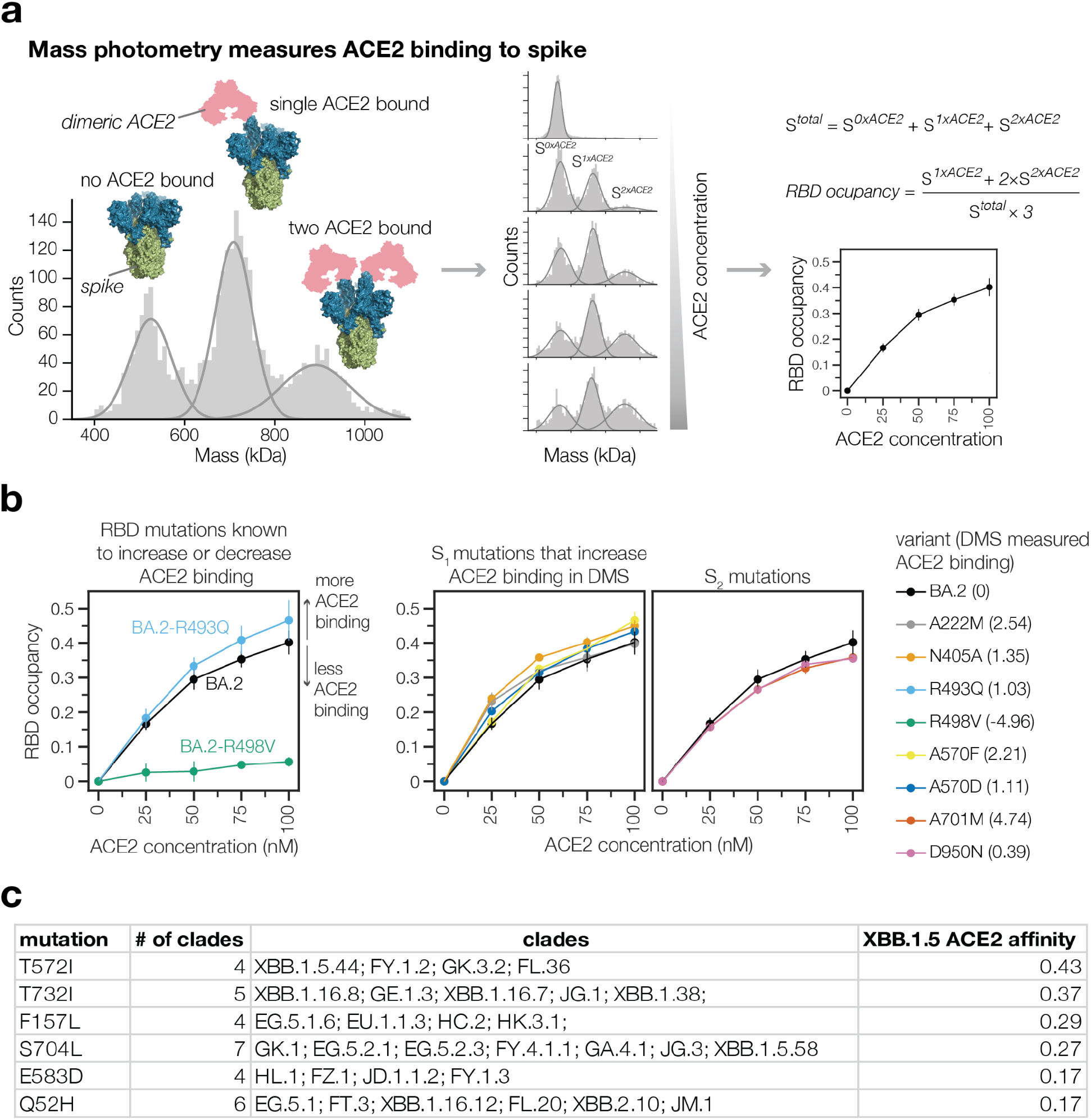
Non-RBD mutations impact ACE2 binding. **a,** ACE2 binding measurements using mass photometry. Histogram on the left shows distribution of spike molecular mass when no (S*^0xACE2^*) one (S*^1xACE2^*) or two (S*^2xACE2^*) ACE2 molecules are bound. We measure how this mass distribution changes as spike is incubated with increasing concentrations of soluble dimeric ACE2. RBD occupancy is the fraction of RBDs bound to ACE2, calculated using Gaussian components for S*^0xACE^*, S*^1xACE2^* and S*^2xACE2^* at each ACE2 concentration. **b,** RBD occupancy measured using mass photometry for different BA.2 spike variants. Left panel shows that a BA.2 spike mutation known to increase ACE2 binding (R493Q/blue) has greater RBD occupancy relative to unmutated BA.2 (black) spike, while a mutation known to decrease ACE2 binding (R498V/green) has lower RBD occupancy. Panels on the right show RBD occupancy for BA.2 spike variants with mutations in S*^1xACE2^*or S*^2xACE2^* subunits measured to increase ACE2 binding in the deep mutational scanning. **c,** Non-RBD mutations measured to increase ACE2 binding in deep mutational scanning experiments that are observed to have arisen independently as defining mutations in at least three XBB-descended clades.

Non-RBD mutations that enhance ACE2 binding have played an important role in SARS-CoV-2 evolution. The following non-RBD mutations that enhance ACE2 binding occurred in major pre-Omicron variants of concern: A570D (Alpha), A222V (several moderate-frequency Delta sublineages), T1027I (Gamma), and D950N (Delta) (**Extended Data Fig. 2d**). In addition, the following non-RBD mutations that occurred in Omicron variants, all of which represent reversions to pre-Omicron residue identities, increase ACE2 binding: K969N, K764N and Y655H. Consistent with prior studies showing that the original D614G mutation increased the proportion of RBDs in the up conformation^21,29,30^, we find that G614D decreases full spike ACE2 binding (**Extended Data Fig. 2d**).

To systematically examine the recent evolutionary role of non-RBD ACE2 binding-enhancing mutations, we tabulated non-RBD mutations that enhance binding and are new mutations in at least three XBB-descended Pango clades (**Fig. 3c**). Some of these mutations may explain why certain clades had a growth advantage. For example, the NTD mutation Q52H provided the EG.5.1 lineage with a clear growth advantage over EG.5^31^, despite not measurably affecting serum neutralization^32^. Our deep mutational scanning provides an explanation for the success of EG.5.1 by showing that Q52H enhances ACE2 binding. Overall, these results suggest that non-RBD mutations that affect ACE2 binding play an important role in SARS-CoV-2 evolution.

### Mapping escape from XBB* infection sera reveals heterogeneity among individuals

We next mapped how mutations in spike affect neutralization by the polyclonal antibodies in sera from 10 vaccinated individuals who either had a confirmed XBB* infection or whose last infection was during a period when XBB lineages were the dominant circulating variants (**Supplementary Table 1**). We performed these measurements with the full spike XBB.1.5 libraries using 293T cells expressing moderate levels of ACE2 that better capture the activities of non-RBD antibodies^33,34^, although the key sites of escape were mostly similar if we used 293T cells expressing high levels of ACE2 or the RBD-only libraries (**Extended Data Fig. 6**). The sites of greatest serum escape were mainly in the RBD (**Fig. 4a-c** and interactive plot at https://dms-vep.github.io/SARS-CoV-2_XBB.1.5_spike_DMS/htmls/summary_overlaid.html). These sites include 357, 371, 420, the 440-447 loop, 455-456, and 473, as well as a few sites in the NTD like 200 and 234. At some sites, the escape mutations are strongly deleterious to ACE2 binding (**Fig. 4c**). For instance, mutations at Y473 cause strong neutralization escape but greatly reduce ACE2 binding, likely explaining why they are not common among circulating SARS-CoV-2 variants. In addition, only some of the antibody-escape mutations mapped in our experiments are accessible by single-nucleotide mutations to XBB.1.5 (**Fig. 4c**). Several escape mutations that are single-nucleotide accessible and do not strongly impair ACE2 binding are found in recent variants, including mutations at site 456 in EG.5.1 and many other XBB variants, mutations at 455 in HK.3.1 and JN.1, mutations at 420 in GL.1, and mutations at 200 in XBB.1.22^7,31^.

**Fig. 4:**
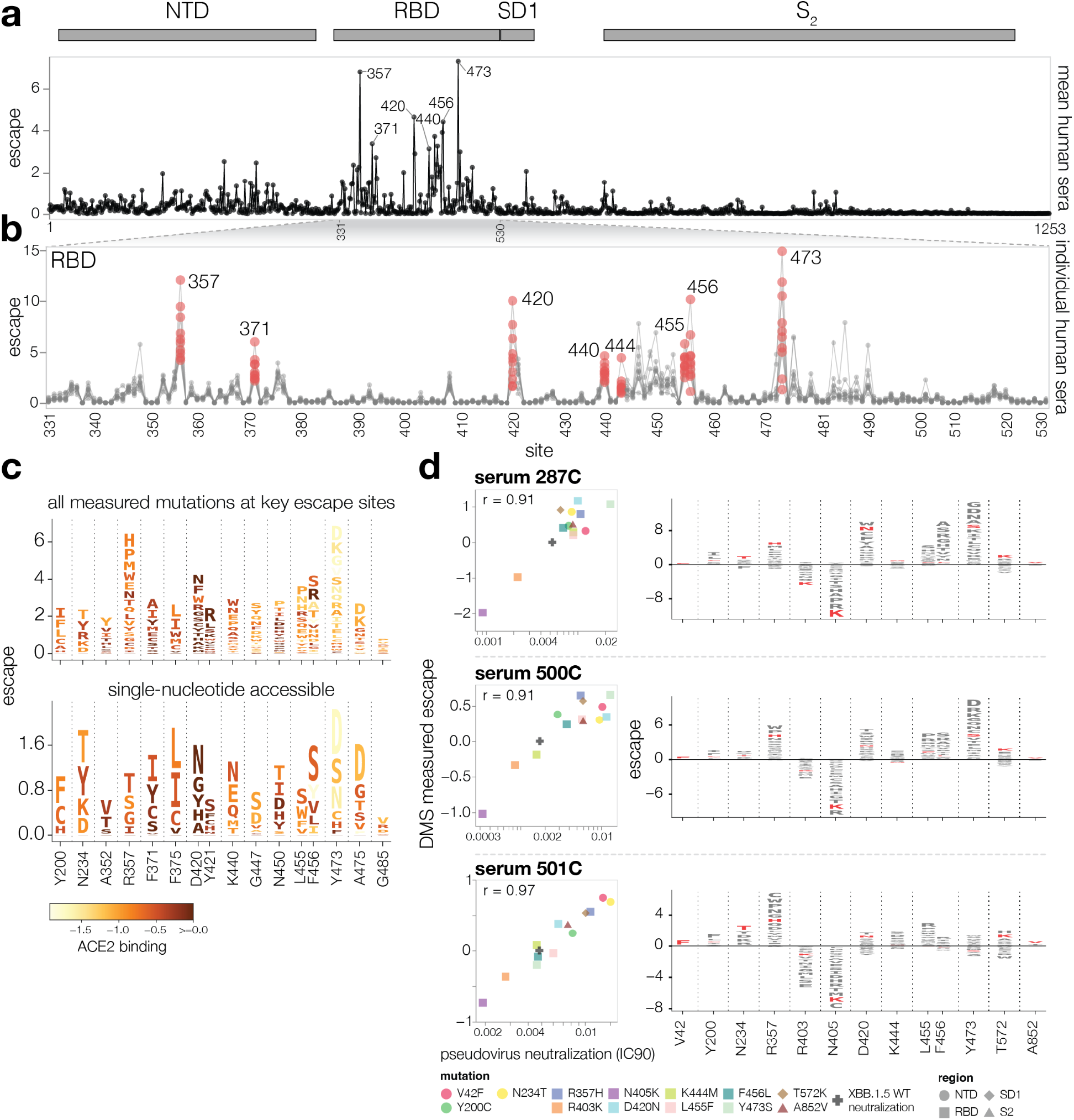
Serum antibody escape mutations for individuals with prior XBB* infections. **a,** Escape at each site in the XBB.1.5 spike averaged across 10 sera collected from individuals with prior XBB* infections. The points indicate the total positive escape caused by all mutations at each site. See https://dms-vep.github.io/SARS-CoV-2_XBB.1.5_spike_DMS/htmls/summary_overlaid.html for an interactive version of this plot with additional mutation-level data. **b,** Zoomed view of the escape at each site in RBD with each line representing one of the 10 sera. Key sites are labeled with red circles indicating escape for each of the 10 sera. **c,** Logo plots showing the 16 sites of greatest total escape after averaging across the sera. Letter heights indicate escape caused by mutation to that amino acid, and letters are colored light yellow to dark brown depending on the impact of that mutation on ACE2 binding (cf. color key). The top plot shows all amino-acid mutations measured, and the bottom plot shows just amino acids accessible by a single nucleotide mutation to the XBB.1.5 spike. **d,** Left: correlation between DMS escape scores and pseudovirus neutralization assay IC90 values for three sera. Right: logo plot showing escape for all sites with mutations validated in the neutralization assays, with the specific validated mutations in red.

While the same mutations often escape many sera, there is also heterogeneity such that the sera-average is not fully reflective of the impacts of mutations on any individual serum (**Fig. 4b,d** and **Extended Data Fig. 7**). For example, while mutations to site Y473 strongly escape neutralization by most sera, two sera we analyzed (493C and 501C) are largely unaffected by mutations at that site. Other key sites of escape, including 420 and 456, show similar heterogeneity across sera. To validate that escape mutations can have very different effects across sera, we performed standard pseudovirus neutralization assays^5^ against a panel of point mutants to the XBB.1.5 spike (**Fig. 4d**). The changes in neutralization in these validation assays were highly correlated with the escape measured by deep mutational scanning, and confirmed the serum-to-serum heterogeneity. For example, Y473S strongly reduces neutralization by sera 287C and 500C, but actually slightly increases neutralization by serum 501C. Similarly, F456L substantially reduces neutralization by only some sera (**Fig. 4d**).

The deep mutational scanning identifies mutations that increase as well as escape neutralization (**Extended Data Fig. 8**). Sensitizing mutations often occur at sites that are mutated in XBB.1.5 relative to earlier variants, such as sites 373, 405, 417, 460, 486 and 505 (**Extended Data Fig. 8**). Presumably in many cases, reverting mutations at these sites restores neutralization by antibodies elicited by infection or vaccination with earlier viral strains. To confirm that the sensitizing mutations identified in the deep mutational scanning actually increased neutralization, we validated the sensitizing effects of R403K and N405K in standard pseudovirus neutralization assay (**Fig. 4d**).

### Some mutations that strongly affect neutralization modulate RBD conformation rather than directly affecting antibody binding

Most sites of strong escape described in the previous section are proximal to the ACE2 binding motif in the RBD that is the target of many potent neutralizing antibodies^35,36^ (we define ACE2-proximal RBD residues as those within 15 Å of ACE2 in the RBD-ACE2 crystal structure). However, the deep mutational scanning also reveals individual mutations at non-RBD or ACE2-distal RBD sites that strongly escape neutralization. Some of these sites, such as 42, 200, 234 in the NTD, 572 in SD1, and 852 in S2 have mutations that cause as much escape as ACE2-proximal RBD mutations, decreasing serum neutralization by as much as 6-fold (**Fig. 4d**). But whereas most mutations at any given site have similar effects on escape at many ACE2-proximal RBD sites, different mutations at the same non-RBD or ACE2-distal RBD site can have diverse effects on neutralization (**Fig. 5a-c**). Furthermore, there is a strong correlation between mutational effects on neutralization and ACE2 binding at these sites: mutations that reduce neutralization also reduce ACE2 binding, and mutations that increase neutralization also increase ACE2 binding (**Fig. 5a,b**). No such consistent correlation exists between neutralization and ACE2 binding for RBD escape sites in close proximity of ACE2 binding interface (**Fig. 5c**).

**Fig. 5:**
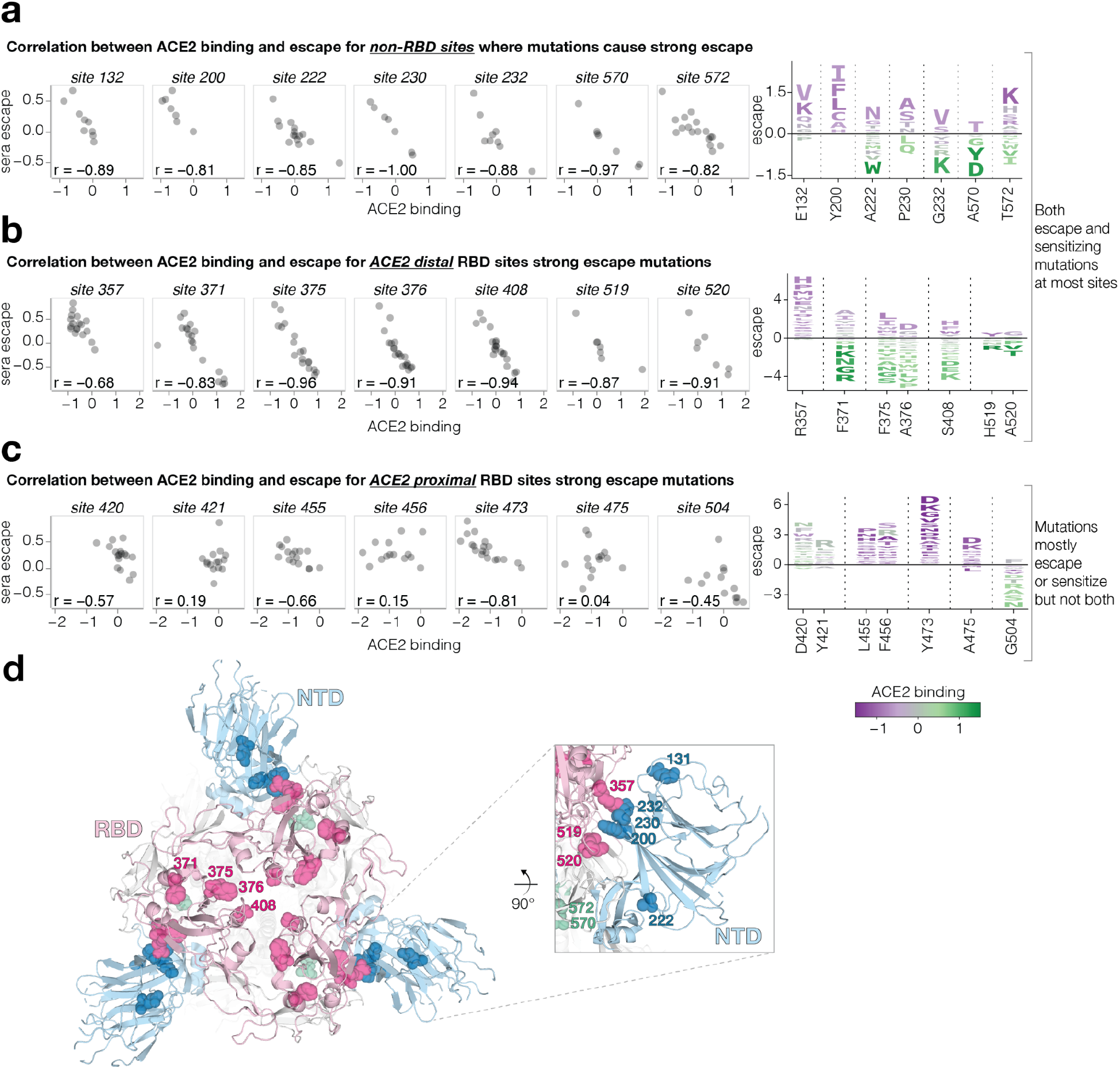
Sera escape and ACE2 binding are inversely correlated for non-RBD and ACE2-distal RBD sites. **a,** Left: correlation between ACE2 binding and escape for the non-RBD sites with the highest mutation-level sera escape. Right: logo plot for the same sites, with letter heights proportional to escape (negative heights mean more neutralization), and letter colors indicating effect on ACE2 binding (green means better binding). **b,** A similar plot for RBD sites that are distal (at least 15 Å) from ACE2. **c,** A similar plot for RBD sites proximal to ACE2. Only sites with at least seven different mutations measured are included in the logo plots. **d,** Top-down view of XBB spike (PDB ID: 8IOT) with the non-RBD and ACE2-distal sites shown in panels a and b highlighted as spheres. The RBD is pink, the NTD is blue, and sites in SD1 are green.

We hypothesize that non-RBD and ACE2-distal RBD mutations that increase both neutralization and ACE2 binding do so by shifting the RBD to a more “up” position, whereas those that decrease neutralization and ACE2 binding do so by shifting the RBD to a more “down” position^37–39^. Indeed, prior work has suggested that mutations that put the RBD in a down position reduce neutralization by class 1 and 4 antibodies that target RBD residues only accessible in the up position, while mutations that put the RBD in a more up position have the opposite effect^14,40^. Our results show that mutations that affect neutralization and ACE2 binding by modulating RBD conformation are common in certain regions of spike—a result that makes structural sense, since most of these mutations are located near the interfaces between the RBD and other spike domains (**Fig. 5d**, **Extended Data Fig. 9**). Furthermore, many of these strong escape sites, including N234, F371, P373, F375, A376, S408, A570, T572, have been previously shown by structural methods to affect RBD conformation^22,37–39,41–44^ or the conformation of key RBD epitopes^19,45^.

### Experimentally measured spike phenotypes partially predict evolutionary success of human SARS-CoV-2 clades

SARS-CoV-2 evolution in humans is characterized by the repeated emergence of new viral clades, which often possess additional amino-acid mutations in spike relative to their predecessors (**Extended Data Fig. 10a**). We estimated the relative growth rates in humans of sufficiently-sampled XBB-descended clades using multinomial logistic regression^46^. As expected, more recent clades generally had higher growth rates, consistent with evolution selecting for viral clades that are more fit (**Fig. 6a**), presumably in part due to the additional mutations in spike^47^.

**Fig. 6:**
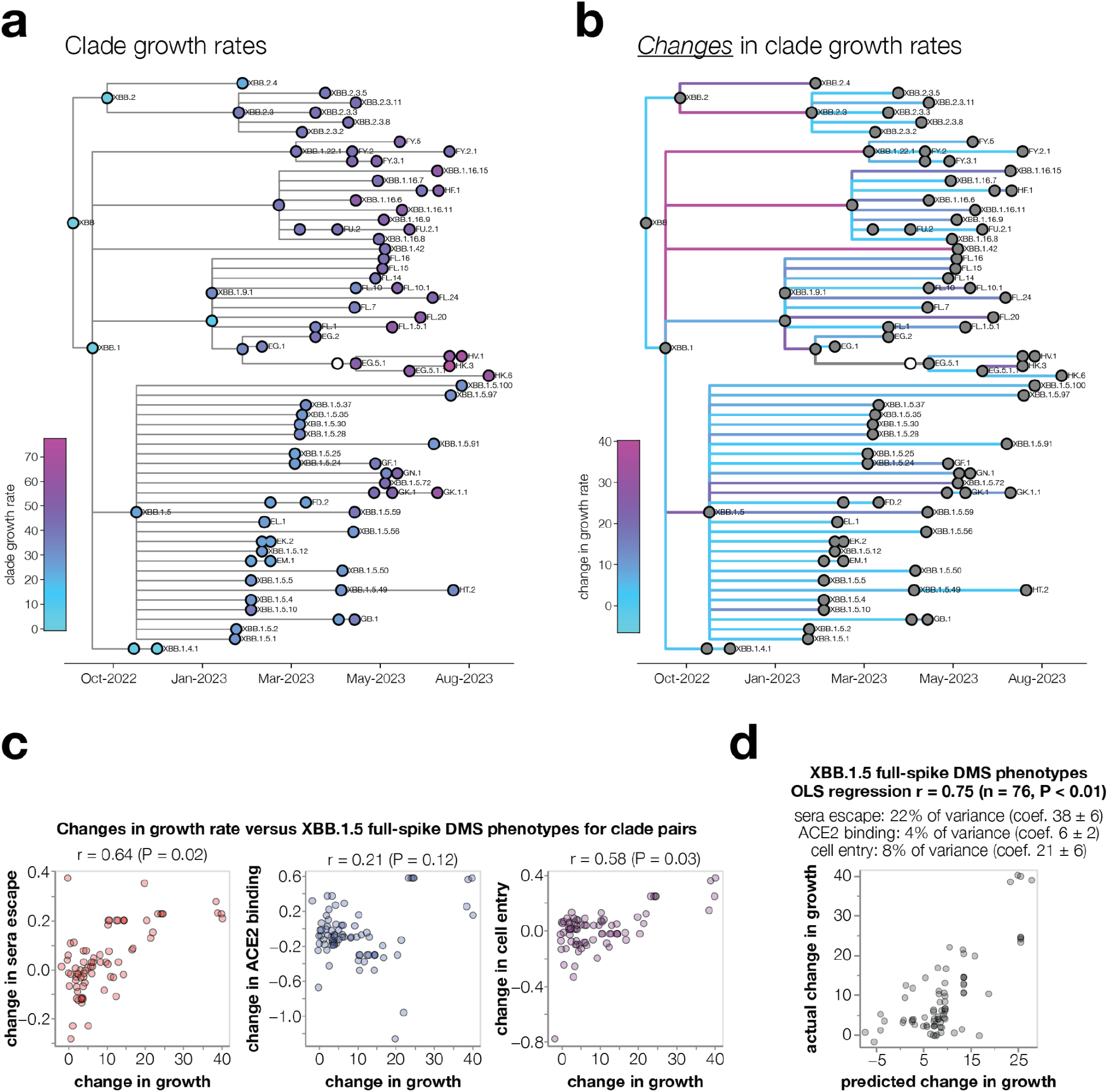
Spike phenotypes measured by deep mutational scanning partially predict the evolutionary success of SARS-CoV-2 clades. **a**, Phylogenetic tree of XBB-descended Pango clades, colored by their relative growth rates. The tree shows only clades with at least 400 sequences and at least one new spike mutation, and their ancestors. Ancestor clades with insufficient sequences for growth rate estimates are in white. **b**, The same phylogeny but with branches colored by the change in growth rate between parent-descendant clade pairs. **c**, Correlation between the changes in growth rate for parent-descendant clade pairs versus the change in each spike phenotype measured in the XBB.1.5 full-spike deep mutational scanning (multiple mutations are assumed to have additive effects). The text above each plot shows the Pearson correlation (r) and a P-value computed by comparing the actual correlation to that for 100 randomizations of the experimental data among mutations. **d**, Ordinary least squares multiple linear regression of changes in growth rate versus all three measured spike phenotypes. The small text indicates the unique variance explained by each variable as well as the coefficients in the regression. See https://dms-vep.github.io/SARS-CoV-2_XBB.1.5_spike_DMS/htmls/current_dms_clade_pair_growth.html and https://dms-vep.github.io/SARS-CoV-2_XBB.1.5_spike_DMS/htmls/current_dms_ols_clade_pair_growth.html for interactive versions of panels **c** and **d** where points can be moused over for details on clades and their mutations.

We sought to determine if the evolutionary success of clades could be predicted from how their mutations affect the spike phenotypes measured by deep mutational scanning. Note that almost any mutation-based measurement (such as just counting mutations) trivially correlates with clade growth because newer clades typically have both better growth rates and more spike mutations (**Fig 6a** and **Extended Data Fig. 10a**). For instance, clade growth rates strongly correlate with the number of spike mutations relative to the early Wuhan-Hu-1 sequence (**Extended Data Fig. 10b**). But this correlation is not informative since the question of evolutionary interest is not whether SARS-CoV-2’s spike will acquire more mutations over time (we already know this will happen), but rather *which* of the various mutant viruses present at any given time will spread. Furthermore, phylogenetic correlations can exaggerate associations between mutations and clade growth^48^. Therefore, we focused our analysis on predicting *changes* in clade growth for each pair of parent-descendant clades separated by at least one spike mutation (**Fig. 6b**). This approach eliminates the confounding effects of phylogenetic relatedness and the accumulation of mutations over time (**Extended Data Fig. 10b,c**), and better answers the question of real evolutionary interest.

Changes in growth between parent-descendant clade pairs were positively correlated with all three experimentally measured spike phenotypes (**Fig. 6c**), with the correlations statistically significant for sera escape and cell entry as assessed by randomization of the measurements among mutations. However, these univariate correlations do not fully capture the information in the experiments, since the effects of mutations on the spike phenotypes are themselves correlated (e.g., mutations that cause sera escape sometimes decrease ACE2 binding). We therefore performed ordinary-least squares multiple linear regression of changes in clade growth versus all three phenotypes. The predictions of this regression correlated with changes in clade growth with a Pearson correlation of 0.75, and were highly statistically significant as assessed by randomization of the measurements among mutations (**Fig. 6d**). Sera escape uniquely explained the largest fraction of the variance in changes in clade growth, but ACE2 binding and cell entry effects also explained some variance. In contrast, neither RBD yeast-display deep mutational scanning of antibody escape^9,49^ and ACE2 affinity^20^ nor the EVEscape deep learning model^50^ were better than randomized data at predicting changes in clade growth at a significance level of P = 0.05 (**Extended Data Figs. 10e,f and 11**). Overall, these results show that full-spike deep mutational scanning can partially predict the evolutionary success of human SARS-CoV-2 clades, and that its predictive power exceeds that of several other methods.

## Discussion

Over 16-million human SARS-CoV-2 genomes have been sequenced to date, enabling rapid identification of variants with new mutations at the sequence level. However, interpreting the consequences of these mutations on viral spread in a partially immune population remains a major challenge. Here we show how pseudovirus-based deep mutational scanning can characterize the effects of mutations throughout spike on three distinct phenotypes critical to viral fitness: cell entry, ACE2 binding, and serum antibody escape.

The full-spike pseudovirus data we generate enables several key insights that were not apparent from prior yeast-display RBD deep mutational scanning approaches^2,18,49^. Most obviously, the data encompass all spike domains, not just the RBD. Strikingly, these data show that non-RBD mutations can affect ACE2 binding, probably by altering the conformation of the RBD in the context of the spike trimer (e.g., in up versus down position). Such mutations are highly relevant for SARS-CoV-2 evolution – for instance, enhancement of ACE2 binding by non-RBD mutations appears to explain why EG.5.1 spread so rapidly after it acquired Q52H, why A222V subvariants of Delta spread widely, and why A570D was selected in Alpha.

Pseudovirus deep mutational scanning also enables us to directly measure how mutations affect neutralization by polyclonal sera. In contrast, prior RBD-display deep mutational scanning could only measure how mutations affect antibody binding^2^, and so to estimate mutational effects on serum neutralization escape it was necessary to characterize hundreds of individual antibodies assumed to represent the polyclonal neutralizing repertoire of humans^3,9^. The ability to directly map how mutations affect serum neutralization leads to two new insights. First, it reveals the heterogeneity in how mutations affect neutralization by sera from different individuals. For instance, we characterize sera from XBB* infected individuals that are both strongly affected and almost completely unaffected by mutations at key sites like 456 or 473. This person-to-person heterogeneity in the antigenic effects of spike mutations will increase as individuals accumulate increasingly distinct exposure histories, and could come to play an important role in shaping SARS-CoV-2 evolution and disease susceptibility as it does for influenza virus^51–53^.

The second major insight from direct mapping of serum escape is that mutations outside the RBD can have marked effects on neutralization. For instance, NTD mutations such as Y42F and N234T decrease neutralization by some sera by nearly 6-fold. The existence of such strong non-RBD escape mutations may seem surprising given that most neutralizing activity in human sera come from antibodies that bind the RBD^2,10,33,54^. However, our data suggest that the strongest non-RBD serum escape mutations act primarily by shifting the RBD to the down conformation, thereby indirectly escaping class 1 and 4 antibodies that bind to RBD surfaces only accessible in the up conformation^14,40^. Of course, such mutations come at a cost to ACE2 binding, since the RBD cannot bind receptor in the down conformation^55,56^. Nonetheless, the ubiquity of such mutations suggests that this mechanism of escape merits monitoring and is in line with prior observations made with endemic human coronaviruses^57,58^. For instance, the RBD of the CoV-229E spike has never been observed in the up conformation^59,60^ despite the fact that this spike somehow manages to bind its receptor during infection. Whether SARS-CoV-2’s spike could eventually evolve to also much more strongly favor a down RBD conformation is unknown.

The most important indication of the relevance of our work is that our measurements of spike phenotypes partially explain the evolutionary success of different SARS-CoV-2 clades in humans. A longstanding goal of evolutionary biology is to understand the molecular phenotypes that contribute to fitness^61^, and then measure them with sufficient accuracy to predict which mutants will actually spread in the real world. We have taken a real step towards this goal, since a simple linear model of the three spike phenotypes measured by our deep mutational scanning explains a substantial amount of the changes in growth rates of XBB-descended clades over the last year. Of course, pseudovirus spike deep mutational scanning will never perfectly predict SARS-CoV-2 evolution: evolution itself is partially stochastic^62^, pseudovirus experiments do not capture all phenotypes of spike relevant to transmission or multicycle replication, and our experiments completely ignore mutations to non-spike genes that contribute fitness^13,63^. Furthermore, it remains technically challenging for deep mutational scanning to account for epistatic interactions among mutations^64^, and we need modeling approaches that better account for how person-to-person heterogeneity in immune-escape mutations shape viral evolution^51^. However, the fact that our deep mutational scanning has substantial power to explain clade growth shows that we have reached the point where experiments can enable useful predictions about SARS-CoV-2 evolution.

## Methods

### Data accessibility and computer code

The data described in this paper are available in both interactive and numerical form at various levels of detail. For easy interactive visualization of the data, we suggest the following interactive charts of how mutations affect all measured phenotypes after applying a reasonable set of filters to remove lower-confidence measurements:

- XBB.1.5 spike: https://dms-vep.github.io/SARS-CoV-2_XBB.1.5_spike_DMS/htmls/summary_overlaid.html
- BA.2 spike: https://dms-vep.github.io/SARS-CoV-2_Omicron_BA.2_spike_ACE2_binding/htmls/summary_overlaid.html
- XBB.1.5 RBD: https://dms-vep.github.io/SARS-CoV-2_XBB.1.5_RBD_DMS/htmls/summary_overlaid.html For numerical data on mutational effects on all measured phenotypes after applying the same reasonable set of filters, see:
- XBB.1.5 spike: https://github.com/dms-vep/SARS-CoV-2_XBB.1.5_spike_DMS/blob/main/results/summaries/summary.csv
- XBB.1.5 spike, per-serum escape: https://github.com/dms-vep/SARS-CoV-2_XBB.1.5_spike_DMS/blob/main/results/summaries/per_antibody_escape.csv
- BA.2 spike: https://github.com/dms-vep/SARS-CoV-2_Omicron_BA.2_spike_ACE2_binding/blob/main/results/summaries/summary.csv
- XBB.1.5 RBD: https://github.com/dms-vep/SARS-CoV-2_XBB.1.5_RBD_DMS/blob/main/results/summaries/summary.csv

In addition to the above interactive charts and numerical data, the entire computational pipelines are available along with rich interactive HTML displays of results. These numerical data and HTML displays include additional options to filter the data for higher and lower confidence values, such as by examining the measurements in each of the two replicate libraries or filtering measurements by how many variants a mutation is seen in. Specifically, full interactive HTML documentation for each deep mutational scanning experiment are rendered on GitHub Pages at::

- XBB.1.5 full spike: https://dms-vep.github.io/SARS-CoV-2_XBB.1.5_spike_DMS/
- BA.2 full spike:https://dms-vep.github.io/SARS-CoV-2_Omicron_BA.2_spike_ACE2_binding/
- XBB.1.5 RBD: https://dms-vep.github.io/SARS-CoV-2_XBB.1.5_RBD_DMS/ GitHub repositories with the actual computer code as well as numerical data are at:
- XBB.1.5 spike: https://github.com/dms-vep/SARS-CoV-2_XBB.1.5_spike_DMS
- BA.2 spike: https://github.com/dms-vep/SARS-CoV-2_Omicron_BA.2_spike_ACE2_binding
- XBB.1.5 RBD: https://github.com/dms-vep/SARS-CoV-2_XBB.1.5_RBD_DMS

Note that most of the analysis in these GitHub repos is performed using dms-vep-pipeline-3 (https://github.com/dms-vep/dms-vep-pipeline-3), version 3.5.3.

Raw sequencing data files have been uploaded under BioProjects: PRJNA1034580 for XBB.1.5 full spike library, PRJNA1035795 for XBB.1.5 RBD-only library, PRJNA1035933 for BA.2 full spike library.

### Design of deep mutational scanning libraries

Deep mutational scanning libraries were designed with codon-optimized XBB.1.5 and BA.2 spikes. The sequence of the XBB.1.5 spike is at https://github.com/jbloomlab/SARS-CoV-2-XBB.1.5_Spike_DMS_validations/blob/main/plasmid_maps/3779_pH2rU3_ForInd_XBB15_Sinobiological_CMV_ZsGT2APurR.gb and the BA.2 spike is at https://github.com/jbloomlab/SARS-CoV-2-XBB.1.5_Spike_DMS_validations/blob/main/plasmid_maps/3332_pH2rU3_ForInd_Omicron_sinobiological_BA2_B11529_Spiked21_T7_CMV_ZsGT2APurR.gb. Note that due to an error on our part early in library design, the XBB.1.5 spike used for libraries lacks F490S mutation present in XBB* variants.

The XBB.1.5 full spike libraries were designed to include all accessible and tolerated mutations by including mutations that appeared in more than 50 sequences on GISAID^65^, occurred independently at least 15 times on pre-built SARS-CoV-2 phylogenies from UShER^66^ or occurred independently at least 2 times in any of the following clades: BA.2.75, BQ.1.1, XBB, XBB.1.5. Deletions that met the above criteria were only included if they occurred in the NTD and we specifically added deletions at sites 342-349, 444-449, and 483-486. We also performed saturating mutagenesis on the sites that met the following criteria: occurred independently at least 2500 times on pre-built SARS-CoV-2 phylogenies from UShER or occurred independently at least 100 times in the clades mentioned above. We also saturated mutations at sites that had strong antigenic effects or otherwise were of special interest^3,9^ full list of these sites can be found at https://github.com/dms-vep/SARS-CoV-2_XBB.1.5_spike_DMS/blob/main/library_design/config.yaml under *sites_to_saturate*. The full list of mutations included in the XBB.1.5 full spike libraries can be found at https://github.com/dms-vep/SARS-CoV-2_XBB.1.5_spike_DMS/blob/main/library_design/results/mutation_design_classification.csv.

For the XBB.1.5 RBD-only libraries, every position in the RBD (positions 331-531) was mutagenized to all possible amino acids.

For the BA.2 full spike libraries the design of mutations to be included in the library was performed the same way as described previously for BA.1 libraries^4^. The final list of mutations in BA.2 libraries can be found at https://github.com/dms-vep/SARS-CoV-2_Omicron_BA.2_spike_DMS/blob/main/library_design/results/aggregated_mutations.csv. Note that the BA.2 libraries used in this study are the same ones briefly described in Haddox at al. 2023^11^.

Analysis pipelines for designing mutagenesis primers are provided at https://github.com/dms-vep/SARS-CoV-2_XBB.1.5_spike_DMS/tree/main/library_design for XBB.1.5 full spike libraries, at https://github.com/dms-vep/SARS-CoV-2_XBB.1.5_RBD_DMS/tree/main/library_design for XBB.1.5 RBD-only libraries, and at https://github.com/dms-vep/SARS-CoV-2_Omicron_BA.2_spike_DMS/tree/main/library_design for BA.2 libraries.

### Production of plasmid libraries used to generate deep mutational scanning libraries

Libraries of lentivirus backbone plasmids containing mutagenised XBB.1.5 or BA.2 spikes were made as described previously^4^. In brief, primers containing desired mutations described above were ordered from IDT as Oligo Pools. Full list of these primers for XBB.1.5 full spike library can be found at https://github.com/dms-vep/SARS-CoV-2_XBB.1.5_spike_DMS/blob/main/library_design/results/oPools.csv, for for XBB.1.5 RBD only library at https://github.com/dms-vep/SARS-CoV-2_XBB.1.5_RBD_DMS/blob/main/library_design/results/oPools.csv, and for BA.2 library at https://github.com/dms-vep/SARS-CoV-2_Omicron_BA.2_spike_ACE2_binding/tree/main/library_design/results (see csv files ending in *oPool.csv*). These primers were used to mutagenize spike sequences using PCR that involves multiple rounds of PCR mutagenesis reactions^67^. Number of PCR rounds and cycles determines the number of mutations per spike introduced and we targeted ∼2-3 mutations per spike, although the precise number of mutations per spike is determined only after lentiviral genomes have been integrated into cells and sequenced with long-read sequencing (see *Long-read PacBio sequencing for variant-barcode linkage* section below). For both XBB.1.5 full spike and RBD-only libraries we pooled spike mutagenesis primers at 2:1 molar ratio between mutations that occur independently multiple times on spike phylogenetic tree and those that occurred multiple times on spike sequences deposited on GISAID database (for RBD only libraries the latter included all possible RBD mutations). For both XBB.1.5 full spike and RBD-only libraries a single round of 10 PCR cycles was used to mutagenize the spike sequence. For BA.2 full spike libraries the same primer pooling strategy and the same number of mutagenesis cycles were used as described for BA.1 libraries^4^. Template spike sequences used for mutagenesis were amplified from https://github.com/jbloomlab/SARS-CoV-2-XBB.1.5_Spike_DMS_validations/blob/main/plasmid_maps/3779_pH2rU3_ForInd_XBB15_Sinobiological_CMV_ZsGT2APurR.gborInd_XBB15_Sinobiological_CMV_ZsGT2APurR.gb plasmid for XBB.1.5 libraries and from https://github.com/jbloomlab/SARS-CoV-2-XBB.1.5_Spike_DMS_validations/blob/main/plasmid_maps/3332_pH2rU3_ForInd_Omicron_sinobiological_BA2_B11529_Spiked21_T7_CMV_ZsGT2APurR.gb plasmid for BA.2 libraries. Spikes for both variants were amplified using *VEP_amp_for* (5′CAGCCGAGCCACATCGCTC) and 3′rev_lib_LinJoin_KHDC (5′CGGAAGAGCGTCGTGTAGGGAAAG) primers. After mutagenesis reaction spike sequences were barcoded in a PCR reaction using primers that contained a unique 16 nucleotide barcode that adds barcodes downstream of spike STOP codon. All libraries had two biological replicates (Lib-1 and Lib-2), which represent two independently produced libraries where mutations in spike are associated with unique barcodes in unique combinations with other mutations. Mutagenised and barcoded spike sequence templates were then added into MluI and XbaI digested lentivirus backbone (Addgene #204579) using HiFi reaction (NEB E2621L). Ampure XP bead purified HiFi reactions were then electroporated into 10-beta electrocompetent E. coli cells (NEB, C3020K) and plated overnight. At least 10 electroporation reactions were performed for each plasmid library in order to produce > 2 million CFUs per library. High diversity of barcoded genomes is required in the later steps of library production in order to minimize barcode duplication, which may happen during lentivirus recombination. For each library bacterial colonies were scraped from overnight plates, pooled and QIAGEN HiSpeed Plasmid Maxi Kit (Cat. No. 12662) was used to prepare plasmid pools used for virus library production.

### Production of cell stored deep mutational scanning libraries

Steps for producing cell-stored spike deep mutational scanning libraries have been described in detail previously^4^ . In brief, two 6-well plates of 293T cells were transfected with plasmid pools described above, lentivirus helper plasmids (BEI: NR-52517, NR-52519, NR-52518) and VSV-G expression plasmid (Addgene #204156). This produced a VSV-G pseudotyped lentivirus pool carrying mutagenised spike sequences in their genomes. VSV-G pseudotyped viruses were then used to infect 293T-rtTA cells at low multiplicity of infection so that no more than one virus would infect each cell. Reverse tetracycline-controlled transactivator (rtTA) is required to induce expression from inducible TRE3G promoter in the lentivirus backbone in the presence of doxycycline (see Addgene #204579 plasmid structure). Note, that 293T-rtTA cells used here is a specific cell clone we isolated when producing rtTA overexpressing cells, which is especially good at producing high titers virus stocks that are required for successful library production. We described production of these 293T-rtTA cells previously^4^. VSV-G infection step was also used to bottleneck the libraries to the desired number of variants; we aimed for between 50,000 and 100,000 variants per library. Final number of variants in each library is shown in **Extended Data Fig. 1a**. After VSV-G infection, cells with successful lentivirus integration were selected for using puromycin. Puromycin selection was performed until visual inspection showed a pure population of cells express zsGreen (which is part of lentivirus backbone, see plasmid Addgene #204579). At this point all cell stored libraries were frozen until further use.

### Long-read PacBio sequencing for variant-barcode linkage

Analysis of linkage between mutations in lentivirus backbone encoded spikes and the barcodes they are associated with was performed using long read PacBio sequencing as described previously^4^. First, we rescued VSV-G pseudotyped viruses from cell-stored libraries by transfecting those cells with lentivirus helper and VSV-G expression plasmids. VSV-G pseudotyped viruses produced from these libraries were then used to infect 293T cells and nonintegrated viral genomes were recovered as described previously^4^. To avoid strand switching and mixing of variant-barcode pairs viral genomes were then minimally PCR amplified using primers with tags that allow to detect strand switching via sequencing. Long read sequencing was performed with PacBio Sequel IIe machine. Consensus variant-barcode sequence was determined requiring at least two CCS sequences per barcode. Variant-barcode lookup tables for each library can be found at:

- For XBB.1.5 full spike library https://github.com/dms-vep/SARS-CoV-2_XBB.1.5_spike_DMS/blob/main/results/variants/codon_variants.csv
- For XBB.1.5 RBD only https://github.com/dms-vep/SARS-CoV-2_XBB.1.5_RBD_DMS/blob/main/results/variants/codon_variants.csv
- For BA.2 library https://github.com/dms-vep/SARS-CoV-2_Omicron_BA.2_spike_ACE2_binding/blob/main/results/variants/codon_variants.csv

Long read sequencing data was also used to determine the average spike mutation frequency in each library. For XBB.1.5 full spike library Lib-1 and Lib-2 mutation frequency 1.91 mutations per spike. For XBB.1.5 RBD only library Lib-1 had an average of 1.82 mutations per spike and Lib-2 had an average of 1.9 mutations per spike. BA.2 libraries had an average of 2.32 and 2.33 mutations per spike for Lib-1 and Lib-2 libraries, respectively.

### Cell entry effect measurement using deep mutational scanning libraries

Cell entry effects for each variant were measured as described previously^4^. In brief, ∼1.5 million transcription units of spike pseudotyped library viruses and ∼5 million of VSV-G pseudotyped transcription units made from the same cell-stored libraries were used to infect target cells. For spike pseudotyped libraries 293T-cells either overexpressing high amounts of ACE2 (described in Crawford et al 2020^5^) or cell expressing medium amount of ACE2 (described in Farrell et al. 2021^33^) were used. Whenever cells were plated for infection with spike-pseudotyped viruses (including for ACE2 and sera selections described below) cells were additionally supplemented with 2.5 µg/ml of amphotericin B (Sigma, A2942) at the time of plating, which we have previously shown^4^ to increase virus titers. For VSV-G pseudotyped libraries 293T cells were used (we used cells not expressing any ACE2 in order to avoid any selection of spike, which can still be present on the surface of these VSV-G pseudotyped viruses). 12-15 hours post infection unintegrated viral genomes were recovered using QIAprep Spin Miniprep kit and prepared for Illumina sequencing as described previously^4^.

For each variant functional score was calculated by getting a log enrichment ratio: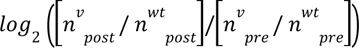, where *n^v^_post_* is the count of variant v in the post-selection (spike-pseudotyped) infection, *n^v^_pre_* is the count of variant v in the pre-selection (VSV-G-pseudotyped) infection, and *n^wt^_post_* and *n^wt^_post_* are the counts of variants without mutations, i.e. wildtype spike, in each condition. Positive functional scores indicate variant is able to enter cells better than wildtype and negative functional scores indicate variant is worse at entering the cells than wildtype. The multi-dms software package^11^ was used to fit a global epistasis model^68^ with a sigmoid global epistasis function to the variant functional scores and to calculate mutation-level effects on cell entry. See https://dms-vep.github.io/SARS-CoV-2_XBB.1.5_spike_DMS/notebooks/func_effects_global_epistasis_Lib1-230614_high_ACE2.html for an example of this fitting for one library; the HTML documentation of the pipeline linked in the *Data availability* section has links to comparable fitting notebooks for each library.

The cell entry effects we describe in the paper are based on the cell entry experiments done on 293T-cells overexpressing high amounts of ACE2 as opposed to medium ACE2 expressing cells. Expression of more ACE2 in 293T-cells leads to higher virus titers on these cells and therefore the fitting of global epistasis model on data from these cells is slightly better.

### Use of non-neutralizable standard for ACE2 binding and serum selection experiments

For both ACE2 binding and serum selection experiments a non-neutralizable standard was used in order to enable conversion of sequence counts to absolute neutralization^4^. We have previously described the use of VSV-G pseudotyped virus as the non-neutralizable standard in antibody selection experiments^4^, and that VSV-G standard was also used for selections with soluble ACE2 protein to measure receptor binding since VSV-G is not neutralized by ACE2. For serum selections, we found that high concentrations of serum appreciably neutralize VSV-G itself making it not suitable as a non-neutralizable standard. We screened multiple alternative viral entry proteins and found that the RDPro glycoprotein, a modified version of an endogenous feline virus RD114 containing HIV R-peptide^69^ that we further modified to contain MLV-A cytoplasmic tail to improve pseudovirus titers^70^, was not neutralized even at high serum concentrations (data not shown). The full sequence of RDPro viral entry protein used in this study can be found at https://github.com/jbloomlab/SARS-CoV-2-XBB.1.5_Spike_DMS_validations/blob/main/plasmid_maps/3737_HDM_RDPro_Twist_MLV-A_HIV-pep_correction.gb. RDPro envelope pseudotyped viruses were produced from cells with integrated barcoded lentivirus genomes as described previously for VSV-G pseudotyped standard^4^. Because RDPro-pseudotyped lentivirus titers were ∼10^4 TU/ml, we further concentrated virus stocks using Lenti-X™ Concentrator (Takara, 631232) to between 3.5*10^5-1.5*10^6 TU/ml.

### Recombinant Protein Production

SARS-CoV-2 spike ectodomain and human ACE2 ectodomains were expressed and purified as described previously^17,19^ Mutant Spike ectodomain constructs were designed in the BA.2 background with HexaPro^28^ mutations, N terminal “MFVFLVLLPLVSS” signal peptide, C terminal GSSG linker, foldon, linker, avi-8x polyhistidine tag, and were placed into pCDNA3.1(+) vector. A222M, N405A, A570F, A570D, A701M, D950N, R493Q, R498V mutations were evaluated in this background. Expi293F cells were diluted to a density of 3 million cells per mL and transfected using ExpiFectamine 293 Transfection Kit (Thermo Fisher Scientific). Cells were incubated shaking at 130 rpm at 37°C and 8% CO_2_. Three to four days post transfection proteins were purified from clarified supernatants. Human ACE2 ectodomains were purified using 1mL Histrap Fast Flowl nickel affinity columns (Cytiva), and washed with 20 mM imidazole, 25 mM sodium phosphate pH 8.0, and 300 mM NaCl prior to elution with an imidazole gradient using a buffer containing 500 mM imidazole, 25 mM sodium phosphate, 300 mM NaCl pH 8.0. SARS-CoV-2 spike ectodomains were purified using 1mL of Ni Excel resin (Cytiva) per purification and washed with 40 mM imidazole, 25 mM sodium phosphate pH 8.0, and 300 mM NaCl prior to elution with 300 mM imidazole, 25 mM sodium phosphate pH 8.0, and 300 mM NaCl. SARS-CoV-2 spike ectodomains were buffer exchanged into 20 mM sodium phosphate pH 8.0, and 100 mM NaCl (PBS) and analyzed by negative stain electron microscopy for proper folding and monodispersity (**Extended Data Fig. 4**). Proteins were concentrated using centrifugal filters (Corning). Human ACE2 ectodomains were further purified by size exclusion chromatography and run through a Superdex 200 Increase 10/300 GL column (Cytiva) pre-equilibrated in PBS. All proteins were analyzed by SDS-PAGE for purity, then flash frozen and stored at −80°C. For deep mutational scanning ACE2 binding experiments biotinylated dimeric ACE2 was purchased from ACROBiosystems (AC2-H82E7-1mg).

### ACE2 binding measurement using deep mutational scanning libraries

Previous research has shown that soluble ACE2 can neutralize SARS-CoV-2 variants with potency proportional to virus binding to the receptor^3,15^. We used this observation to measure the effects of mutations in our deep mutational scanning libraries on ACE2 binding.

As described previously^4^, before starting ACE2 binding experiments we spiked-in a VSV-G non-neutralizable standard at 1-2% of the total virus titers used. ∼1 million virus transcription units per sample were incubated with soluble monomeric or dimeric ACE2 at 37°C for 1 h before being added to 293T-ACE2 cells. 293T-ACE2 cells expressing a medium amount of ACE2 (‘medium-ACE2’ cells described in Farrell et al. 2021^33^) were used for all ACE2 binding experiments. For these experiments we targeted a range of ACE2 concentrations to use that would span from less than IC50 to full virus neutralization in order to capture both mutations that increase ACE2 binding (those that are neutralized by soluble ACE2 very potently) and those that decrease it (which would be more difficult to neutralize with soluble ACE2 neutralized). For monomeric ACE2 the starting concentration was 2.88 µg/ml and it was increased 3-fold for the other dilutions. For dimeric ACE2 starting concentration was 0.21 µg/ml and it was similarly increased 3-fold for the other dilutions. 12-15 hours post infection non-integrated lentiviral genomes were extracted from cells and barcode sequencing libraries were prepared as described previously^4^. ACE2 binding experiments were performed with two biological replicates for each library.

Analysis of mutation-level effects and fitting of neutralization curves to the data was performed using *polyclonal* software^71^ version 6.9. Examples of polyclonal model fitting for monomeric ACE2 data can be found at:

- For XBB.1.5 full spike libraries https://dms-vep.github.io/SARS-CoV-2_XBB.1.5_spike_DMS/notebooks/fit_escape_ACE2_binding_Lib1-230614-monomeric_ACE2.html
- For XBB.1.5 RBD only libraries https://dms-vep.github.io/SARS-CoV-2_XBB.1.5_RBD_DMS/notebooks/fit_escape_ACE2_binding_Lib1-230615-monomeric-ACE2.html
- For BA.2 libraries https://dms-vep.github.io/SARS-CoV-2_Omicron_BA.2_spike_ACE2_binding/notebooks/fit_escape_ACE2_binding_Lib1-230114-monomeric_ACE2.html

The HTML documentation of the pipeline linked in the *Data availability* section has links to comparable fitting notebooks for each replicate library, as well as dimeric ACE2 selection data available for XBB.1.5 RBD only libraries.

### Mass photometry

Mass photometry was performed on a Refeyn TwoMP system (Refeyn Ltd). Microscope cover slides were rinsed with isopropanol and Milli-Q water then dried under nitrogen flow. Sample chambers were assembled using silicon gaskets, instrument lens was coated with immersol and slides were placed on the MP sample stage. Samples were added to the sample chamber and the instrument was focused immediately prior to each data acquisition. Spike ectodomain samples were diluted to 25 nM for all data acquisitions. Spike was mixed with 0, 25, 50, 75 or 100 nM of dimeric human ACE2 ectodomain and incubated at room temperature for 5 min prior to data acquisition. We conducted mass calibration using in-house protein standards. Data was analyzed using DiscoverMP. RBD-ACE2 occupancy was calculated using python script with relative molecule counts from event exports from DiscoverMP. Representative histograms were generated using python script using event exports from DiscoverMP. Mass events were truncated to a range of 0-1100 kDa for spike only runs and to 350-1100 kDa for spike-ACE2 runs. Gaussian mixture models with 3 components and full covariance generated from the Sci Kit Learn Library were overlaid on histogram and latent variables (mean, covariance, weight) for each component were determined. Components were sorted by mean into groups corresponding to molecular masses of 400-600 kDa, 600-800 kDa, 800-1000 kDa for unbound spike, 1 ACE2 bound spike, and 2 ACE2 bound spike, respectively. The relative abundance of each of the three species and the spike-ACE2 occupancy were determined from the respective weight of the Gaussian components, as follows: 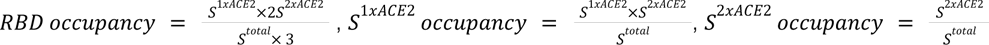, where S*^1xACE2^* is the weight of the respective Gaussian components for single ACE2 bound spike, S*^2xACE2^*is Gaussian component spike bound by 2 ACE2 molecules and S*^total^*is the sum of Gaussian components for spike bound by no, one or 2 ACE2 molecules.

### Serum escape mapping using deep mutational scanning libraries

XBB* infection sera neutralization was determined using standard pseudovirus neutralization assay described in Crawford et al. (2020)^5^. The sequence of the spike expression plasmid used for these experiments is provided at https://github.com/jbloomlab/SARS-CoV-2-XBB.1.5_Spike_DMS_validations/blob/main/plasmid_maps/HDM_XBB15.gb. Using these measurements we determined the amount of serum needed to neutralize the virus at IC98-IC99. As described previously^4^, before starting selections we spiked-in a non-neutralizable standard at 1-2% of the total virus titers used. RDPro pseudotyped non-neutralizable standard was used for all serum selections to avoid non-specific standard neutralization (see section *Use of non-neutralizable for ACE2 and serum selection experiments* above). For sera selection experiments ∼1 million transcription units for each library sample were used. Libraries were incubated at three increasing serum concentrations starting with IC98-IC99 (depending on serum volume available) and increasing serum concentration 4-fold at each dilution. These serum concentrations were selected so that only a small percentage of variants would be able to pass sera selection, therefore selecting for strongest escape variants. Additional sera concentrations are used to cover a greater dynamic range as sometimes neutralization values determined against wild-type spike using luciferase-based system do not quite match neutralization values for library virus pool. Virus-serum mixtures were incubated for 1 h at 37°C and subsequently 293T-ACE2 cells were infected with them. We used medium ACE2 expressing cells for all serum selection experiments (‘medium-ACE2’ cells in Farrell et al. 2021^33^) although as shown in **Extended Data Fig. 6** we did not detect significant differences in serum escape compared to cells with high ACE2 expression. 12-15 hours post infections non-integrated lentiviral genomes were extracted from cells and barcode sequencing libraries were prepared as described previously^4^.

*Polyclonal* software^71^ (version 6.9) was used to analyze mutation-level escape and fit neutralization curves to the data. An example for data fitting for one sera sample can be found at https://dms-vep.github.io/SARS-CoV-2_XBB.1.5_spike_DMS/notebooks/fit_escape_antibody_escape_Lib1-230815-sera-343C_mediumACE2.html for XBB.1.5 full spike library and at https://dms-vep.github.io/SARS-CoV-2_XBB.1.5_RBD_DMS/notebooks/fit_escape_antibody_escape_Lib1-230815-sera-343C_mediumACE2.html for XBB.1.5 RBD library. The HTML documentation of the pipeline linked in the *Data availability* section has links to comparable fitting notebooks for each biological replicate, as well as all other sera fits.

### Validation of escape using standard pseudovirus neutralization assay

To validate serum escape mutations, we cloned desired point mutants into an expression plasmid coding for XBB.1.5 spike. The sequence of this XBB.1.5 expression plasmid is at https://github.com/jbloomlab/SARS-CoV-2-XBB.1.5_Spike_DMS_validations/blob/main/plasmid_maps/3813_HDM_XBB15_with_F490S.gb (note this spike sequence contains F490S mutation). Pseudoviruses were generated and titrated as described in Crawford et al. (2020)^5^ except that pHAGE6_Luciferase_IRES_ZsGreen backbone was used which requires only Gag/Pol (BEI: NR-52517) helper plasmid to produce pseudoviruses. Pseudovirus stocks were diluted to stock concentration of >200,000 relative light units per ul and neutralization assays were performed on medium-ACE2 cells. Starting serum dilution for neutralization assays was 1:30 and it was serially diluted 1:3 to generate neutralization curves. Neutralization curves were plotted by fitting a Hill curve to fraction infectivity data for each variant. This was done using *neutcurve* package (https://jbloomlab.github.io/neutcurve/, version 0.5.7). Analysis notebook for neutralization curves is at https://github.com/jbloomlab/SARS-CoV-2-XBB.1.5_Spike_DMS_validations/tree/main.

### Cells

All cells in this study were maintained in D10 media (Dulbecco’s Modified Eagle Medium with 10% heat-inactivated fetal bovine serum, 2 mM l-glutamine, 100 U/mL penicillin, and 100 μg/mL streptomycin). 293T-ACE2 cells expressing medium amount of ACE2 (‘medium-ACE2’ cells described in Farrell et al. 2021^33^) were additionally supplemented to 2 µg/ml of doxycycline. Cells used to store spike libraries were maintained in media supplemented with doxycycline-free FBS as described previously^4^.

### Ethics statement

XBB* infection sera were collected after informed consent from participants in the prospective longitudinal Hospitalized or Ambulatory Adults with Respiratory Viral Infections (HAARVI) study from Washington State, USA, which was approved by University of Washington Institutional Review Board (protocol #STUDY00000959).

### Comparison of deep mutational scanning phenotypes to changes in clade growth

To estimate clade growth rates, we used a multinomial logistic regression model of global lineage frequency data. GISAID sequences were obtained from the bulk .fasta download (dated 2023-10-02) and processed with Nextclade (v2.14.0) using the BA.2 reference (‘sars-cov-2-21L’). Using Nextclade quality metrics, only sequences with >90% coverage and an overall QC status of "good" were retained. Since outlier dates could distort model estimates, we required sequences to have a fully specified deposition and collection date (ie. year, month, and day), and to have a collection date within 150 days of deposition (primarily to avoid collection dates where the year was incorrectly annotated). Finally, for each lineage, we excluded sequences with dates that were extreme outliers for that specific lineage, falling outside of 3.5 times the interquartile range of the median. For the model fit, we retained countries with >500 sequences, and lineages with >200 sequences. Counts for each lineage were aggregated per country, per day.

Lineage counts were modeled using multinomial logistic regression, with global per-lineage growth rates (ie. shared between all countries), and per-lineage per-country intercepts. The model was implemented in the Julia language and the likelihood maximized using gradient descent, using Flux.jl and CUDA.jl to allow for GPU computation. See https://github.com/MurrellGroup/MultinomialLogisticGrowth/tree/main/plots for visualizations of the model fits.

Growth rates are interpreted such that the ratio of the frequency of two lineages *l_i_* and *l_j_*changes with time as

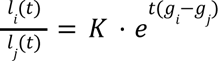

where g_i_ and g_j_ are the growth rates for each lineage, *t* is time in years, and *K* is a constant determined by their intercepts.

The repository estimating the growth rates is at https://github.com/MurrellGroup/MultinomialLogisticGrowth and the actual growth rate estimates are at https://github.com/MurrellGroup/MultinomialLogisticGrowth/blob/main/model_fits/rates.csv. For the analyses in this paper, we considered only growth estimates from XBB descended clades with at least 400 sequences, since clades with more sequences have more accurate growth estimates. The definitions of the clades (e.g., which mutations they contain) as well as their phylogeny (parents and descendants for each clade) were taken from https://github.com/corneliusroemer/pango-sequences/blob/main/data/pango-consensus-sequences_summary.json.

As described in the results and **Fig. 6** and **Extended Data Fig. 10**, directly predicting growth rates of clades from the deep mutational scanning is a confounded approach due to both phylogeny and the simple fact that newer clades tend to have both more spike mutations and higher growth rates, leading to a trivial correlation of clade growth rate with number of spike mutations. The real question of interest is not whether more fit clades with additional mutations will be selected over time (we know they will), but rather which of the mutant clades present at any given time will be more successful. Therefore, as indicated in **Fig. 6b**, we computed the *change* in growth rate for each parent-descendant clade pair with estimates for both clade members and at least one spike mutation. We then also computed the change in each spike phenotype as measured by deep mutational scanning for the clade pairs based on the mutations separating the pair members, simply adding together the mutation effects for pairs separated by multiple spike mutations. Non-spike mutations were ignored. The Pearson correlations with each phenotype are shown in **Fig. 6c**, and the statistical significance of the correlations were assessed by randomizing the deep mutational scanning measurements among mutations 100 times and assessing how many randomizations had a correlation greater than or equal to the observed value. To test the predictive value of combining all spike phenotypes, we performed a similar analysis but using ordinary least squares multiple linear regression on all three phenotypes. Those results are shown in **Fig. 6d**, with the significance again assessed by comparing the actual Pearson correlation to that generated by fitting the model to data randomized among sites. To compute the unique variance explained by each phenotype, we removed the phenotypes one-by-one and computed the unique variance explained as the squared Pearson correlation for the full model minus the squared Pearson correlation for the model with that phenotype removed. See https://dms-vep.github.io/SARS-CoV-2_XBB.1.5_spike_DMS/notebooks/current_dms_compare_natural.html for the notebook performing this analysis, and https://github.com/dms-vep/SARS-CoV-2_XBB.1.5_spike_DMS/blob/main/results/compare_natural/current_dms_clade_pair_growth.csv for the numerical data on the clade pairs and their changes in spike phenotypes.

We compared the predictive value of the full-spike deep mutational scanning to the predictive value of models based on several other values. The first such comparator model simply involves counting the change in number of spike mutations relative to Wuhan-Hu-1 in each clade pair; as shown in **Extended Data Fig. 10c**, this model has no predictive value.

The second comparator model uses the effect of RBD mutations on ACE2 affinity and RBD expression as measured in yeast-display deep mutational scanning of the XBB.1.5 RBD^20^ as well as per-site escape values (same value assigned to each mutation at each site) as computed using the default settings of the antibody escape calculator^9^ at https://github.com/jbloomlab/SARS2-RBD-escape-calc/tree/5ebb88e5b8c9adc1b601b3cb1cc5308532d97a38 which is based on monoclonal antibody deep mutational scanning data^49^. For this model, all non-RBD mutations were assigned a value of zero for all phenotypes. As shown in **Extended Data Fig. 10d,e**, the predictions of this model are not significant at a level of P = 0.05 compared to models with the measurements randomized among sites.

The third comparator model uses the effects of mutations to XBB.1.5 as estimated using the EVEscape method (https://evescape.org/data)^50^. As shown in **Extended Data Fig. 10f**, the predictions of this model are not significant at a level of P = 0.05.

The numerical data used for all the comparator models is at https://github.com/dms-vep/SARS-CoV-2_XBB.1.5_spike_DMS/tree/main/data/compare_natural_datasets.

## Acknowledgments

We thank Ryan Hisner for helpful comments during library design. We thank Cornelius Roemer and Richard Neher for helpful comments on interpreting the effects of mutations in the context of SARS-CoV-2 evolution. We thank Thomas Peacock for useful discussions on ACE2 binding. We gratefully acknowledge all data contributors, including the authors and their originating laboratories responsible for obtaining the specimens, and their submitting laboratories that generated the genetic sequence and metadata and shared via the GISAID Initiative the data on which part of this research is based. JDB is an investigator at the Howard Hughes Medical Institute. D.V. is an Investigator of the Howard Hughes Medical Institute and the Hans Neurath Endowed Chair in Biochemistry at the University of Washington. This work was supported by the NIH/NIAID R01AI141707 grant and contract 75N93021C00015 to J.D.B as well as NIH/NIAID P01AI167966 and DP1AI158186 grants and 75N93022C00036 contract to D.V.), a Pew Biomedical Scholars Award (D.V.), an Investigators in the Pathogenesis of Infectious Disease Awards from the Burroughs Wellcome Fund (D.V.), the University of Washington Arnold and Mabel Beckman cryoEM center and the National Institute of Health grant S10OD032290 (to D.V.). This research was also supported by the Genomics & Bioinformatics Shared Resource, RRID:SCR_022606, of the Fred Hutch/University of Washington Cancer Consortium (P30 CA015704), by the Flow Cytometry Shared Resource, RRID:SCR_022613, of the Fred Hutch/University of Washington/Seattle Children’s Cancer Consortium (P30 CA015704), and by Fred Hutch Scientific Computing, NIH grants S10-OD-020069 and S10-OD-028685. B.M. was funded by SciLifeLab’s Pandemic Laboratory Preparedness programme (VC-2022-0028) and the Erling Persson Foundation (2021 0125).

## Supplementary figures

**Extended Data Fig. 1:**
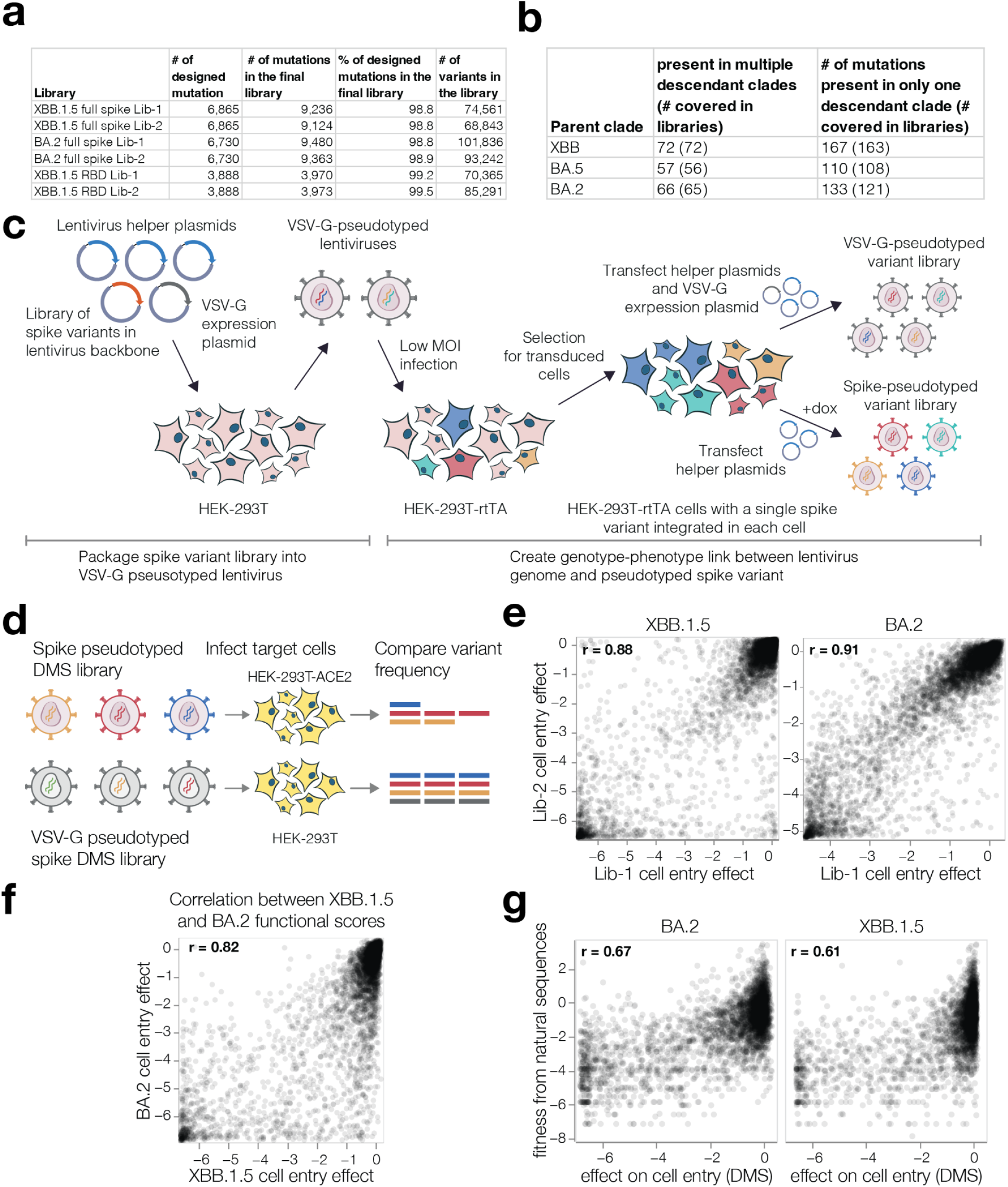
XBB.1.5 and BA.2 spike deep mutational scanning libraries. **a,** Number of targeted and final number of mutations and barcoded variants in the XBB.1.5 and BA.2 full spike and XBB.1.5 RBD pseudovirus-based deep mutational scanning libraries. **b,** Total number of unique mutations present in BA.2 and XBB descendant Pango clades and the number of those mutations that are present in at least three barcoded variants in each replicate of the BA.2 or XBB.1.5 full spike libraries, which was the minimum number of occurrences we needed to make high-confidence estimates of the mutational effects on cell entry. **c,** Method for creating genotype-phenotype linked spike deep mutational scanning libraries, as previously described in Dadonaite et al. (2023)^4^. Lentivirus backbone plasmids encoding barcoded mutagenised spike genes together with helper and VSV-G expression plasmids are transfected into 293T cells to make VSV-G pseudotyped virus. These viruses are used to infect 293T-rtTA cells at MOI < 0.01 so that no more than one spike variant is integrated into each cell. Transduced cells are selected for lentiviral integration, and spike pseudotyped virus libraries are produced from these cells by transfecting helper plasmids in the presence of doxycycline to induce spike expression. In the absence of doxycycline and with added VSV-G expression plasmid, VSV-G pseudotyped virus libraries are also produced from the same cell lines; these VSV-G pseudotyped viruses are used to help estimate effects of spike mutations on cell entry as described in the next panel. **d,** Method used to measure effects of mutations in spike on cell entry. The ability of each spike variant to mediate cell entry is assessed by quantifying its relative frequency in 293T-ACE2 cells infected with spike-pseudotyped versus VSV-G pseudotyped libraries. **e,** Correlations between the effects of mutations on cell entry measured using each of the two independent full spike libraries of XBB.1.5 or BA.2. Throughout the rest of this paper, we report the mean value between the two libraries. **f,** Correlation between mutational effects on cell entry measured for the XBB.1.5 versus BA.2 full spike libraries. **g,** Correlation between mutational effects measured with the XBB.1.5 or BA.2 full spike libraries and fitness effects of those mutations estimated from actual human SARS-CoV-2 sequences^13^.

**Extended Data Fig. 2:**
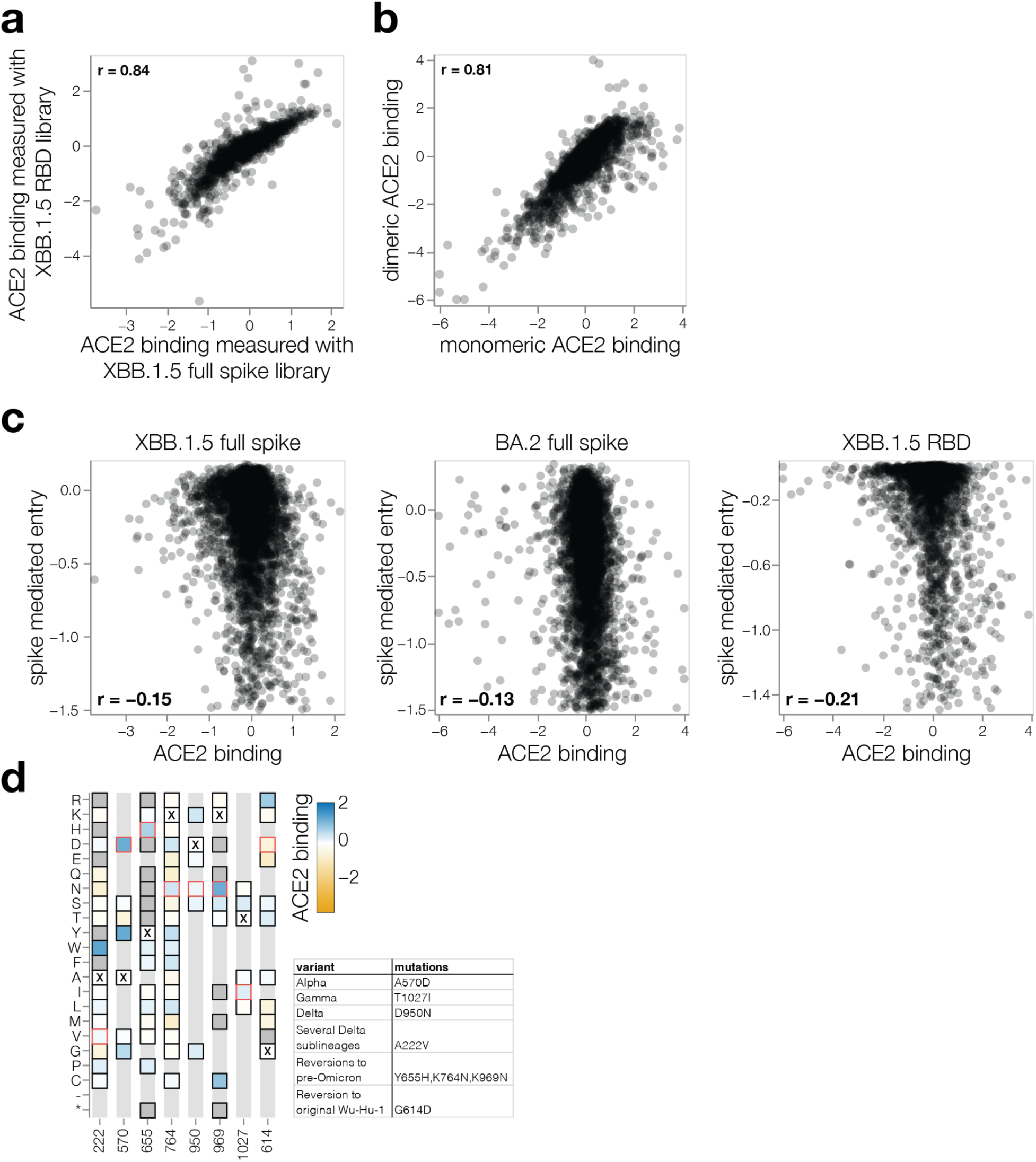
Correlations among measured mutational effects on ACE2 binding. **a,** Correlation between effects of mutations on ACE2 binding measured with XBB.1.5 full spike and XBB.1.5 RBD pseudovirus libraries. **b,** Correlation between effects of mutations on ACE2 binding measured using XBB.1.5 RBD pseudovirus library with monomeric and dimeric ACE2. Heatmaps with the XBB.1.5 RBD pseudovirus measurements made using monomeric and dimeric ACE2 are at https://dms-vep.github.io/SARS-CoV-2_XBB.1.5_RBD_DMS/htmls/monomeric_ACE2_mut_effect.html and https://dms-vep.github.io/SARS-CoV-2_XBB.1.5_RBD_DMS/htmls/dimeric_ACE2_mut_effect.html, respectively **c,** Correlation between effects of mutations on ACE2 binding and spike-mediated cell entry for different libraries. **d,** ACE2 binding heat map showing key sites that have mutated in the past major SARS-CoV-2 variants. Specific variant mutations are highlighted in red outline. Table on the right indicates variants in which these mutations occurred. Interactive plot showing ACE2 binding for all mutations measured in spike is at https://dms-vep.github.io/SARS-CoV-2_XBB.1.5_spike_DMS/htmls/monomeric_ACE2_mut_effect.html.

**Extended Data Fig. 3:**
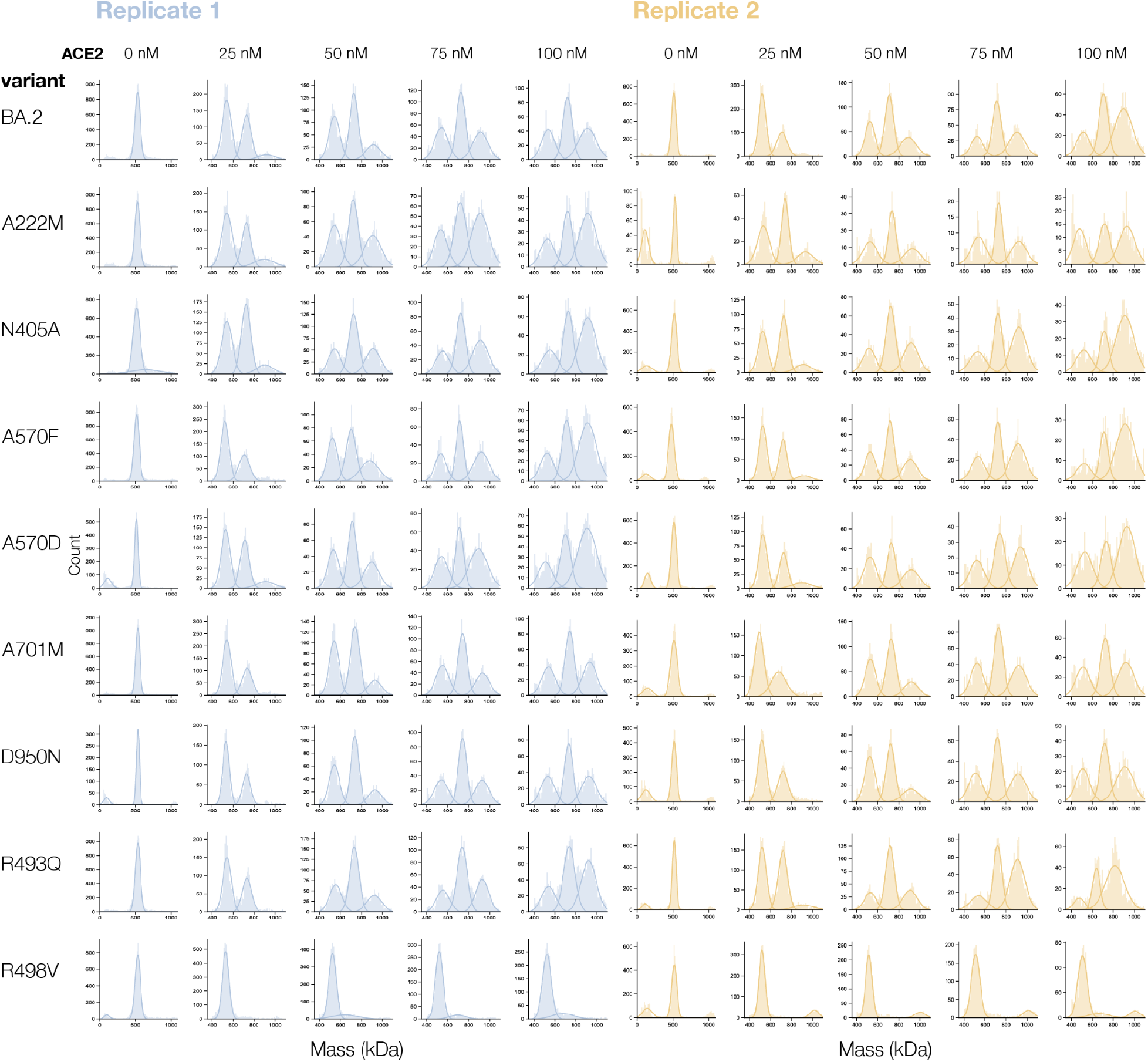
Mass photometry measurements for individual BA.2 spike variants. Spike molecular mass distributions measured using mass photometry for each biological replicate (blue and orange) corresponding to independent purification batches. Each row shows a BA.2 spike mutant and each column shows measurements at different ACE2 concentrations. In the absence of ACE2, some samples had a small peak to the left which may be a misfolded spike monomer^72^ which was present only in some protein preparations and is excluded from Gaussian curve fitting in the presence of ACE2.

**Extended Data Fig. 4:**
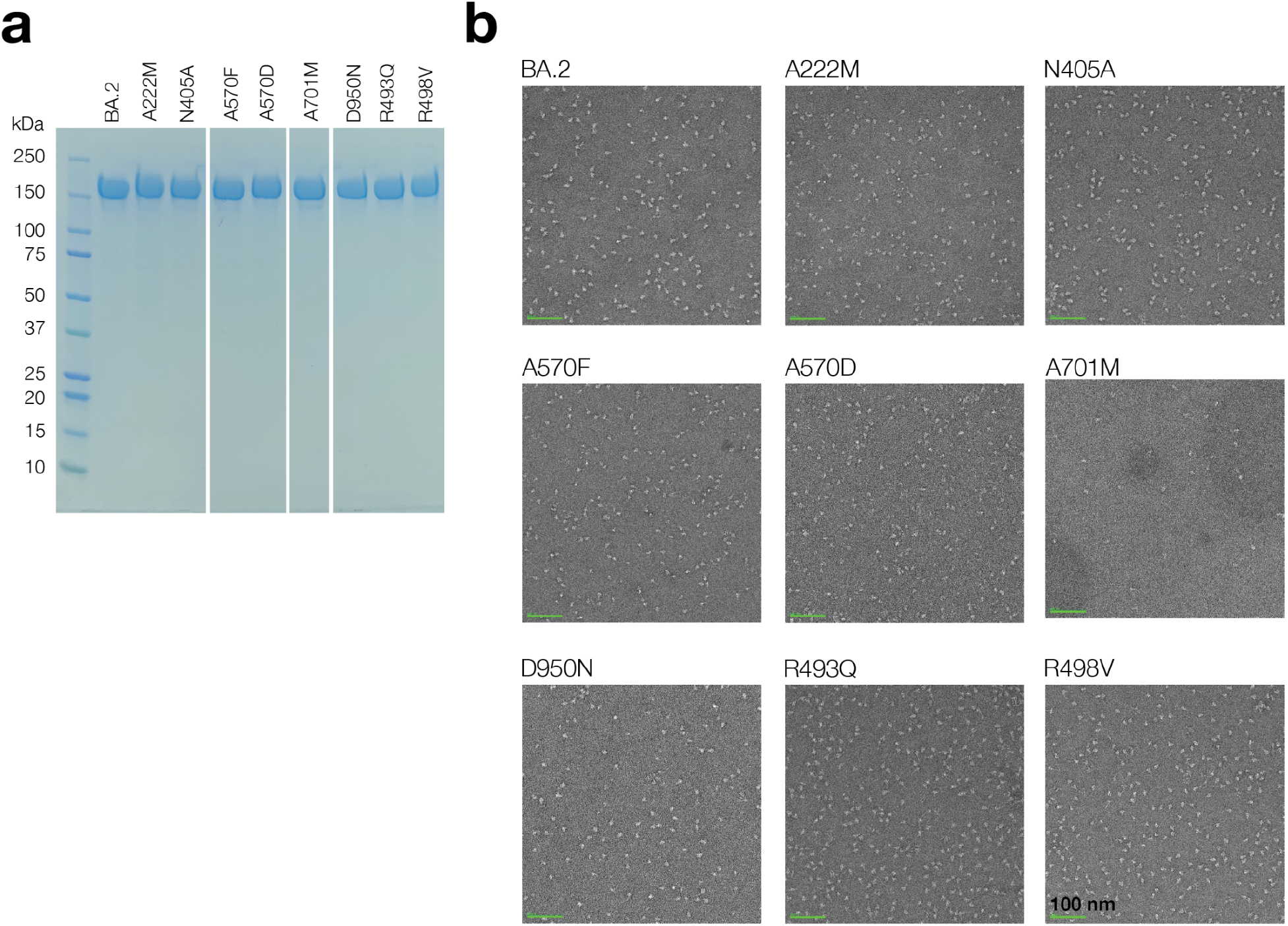
BA.2 and its mutant spike preparations. **a,** Reducing SDS-PAGE gel for purified BA.2 wildtype and mutant spike ectodomains. All constructs are pre-fusion stabilized with HexaPro mutations^28^. 3µg of purified protein loaded. Single major band for all samples confirms sample purity. **b,** Negative stain electron microscopy images for the purified spike mutants to confirm proper folding and monodispersity of the samples.

**Extended Data Fig. 5:**
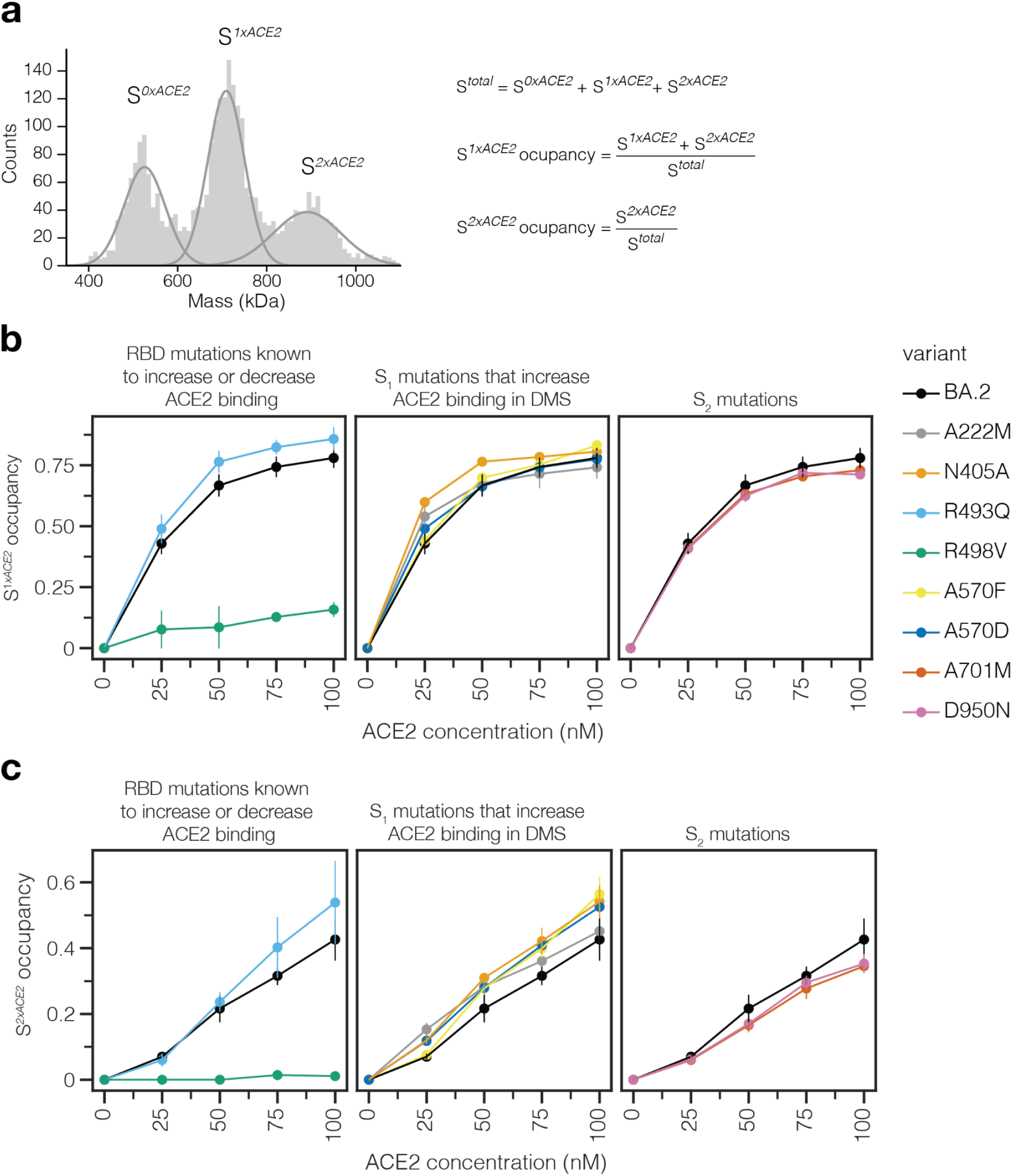
Mass photometry measurements for S_1_ and S_2_ occupancy. **a,** Illustration of Gaussian components for no (S*^0xACE2^*) one (S*^1xACE2^*) or two (S*^2xACE2^*) ACE2-bound spike. S*^1xACE2^* occupancy is the fraction of spikes bound by one ACE2 molecule and S*^2xACE2^* is the fraction of spikes bound by two ACE2 molecules. **b,** S*^1xACE2^*occupancy measured using mass photometry for different BA.2 spike mutants. **c,** S*^2xACE2^* occupancy measured using mass photometry for different BA.2 spike mutants

**Extended Data Fig. 6:**
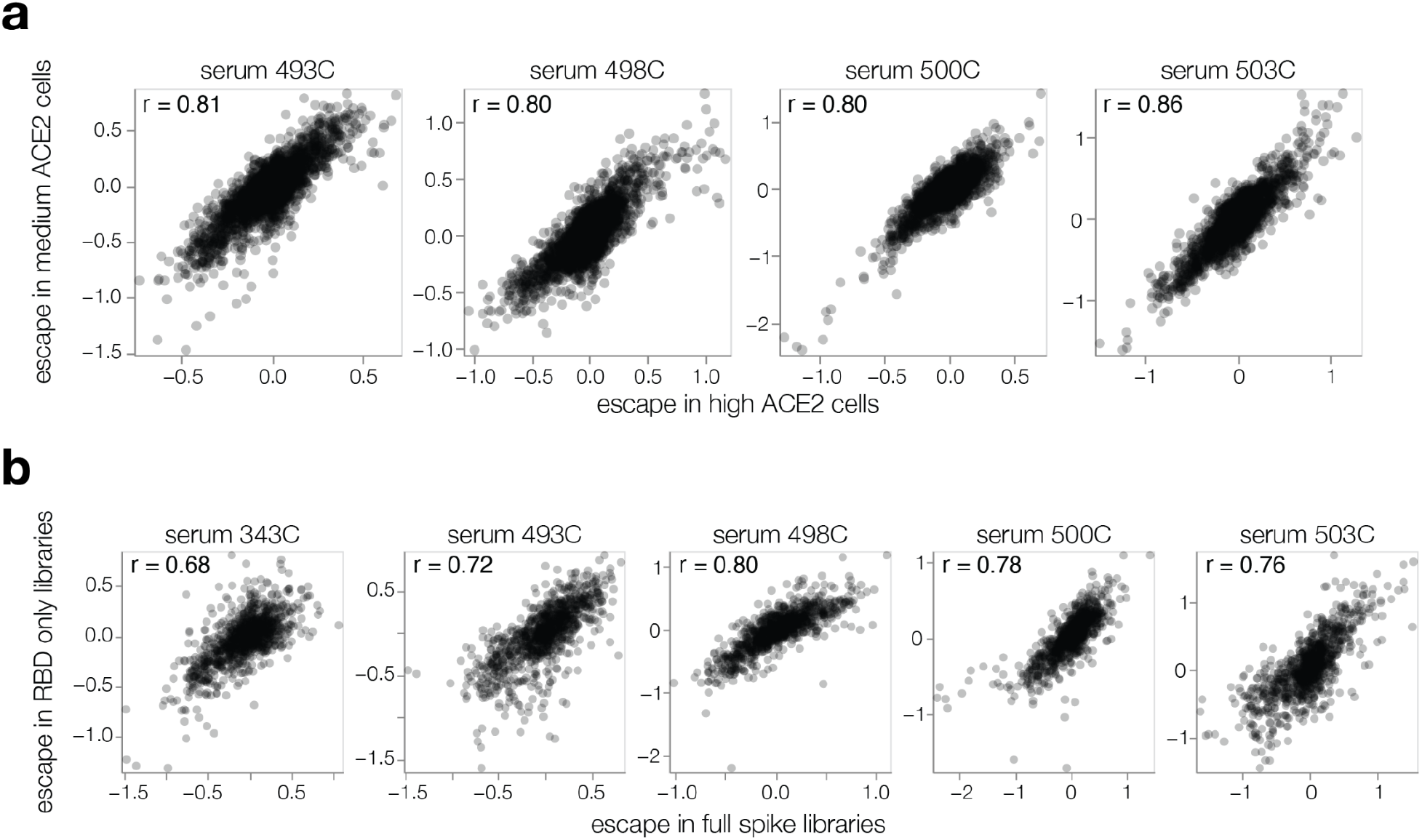
Correlation among serum escape mapping experiments. **a,** Correlation between mutation escape scores for experiments using the full-spike XBB.1.5 libraries performed on 293T cells expressing high or medium amounts of ACE2 for four sera. Note that the medium cells were used for all other figures shown in this paper. **b,** Correlation between mutation escape scores for mutations in the XBB.1.5 full spike and RBD-only libraries. See https://dms-vep.github.io/SARS-CoV-2_XBB.1.5_spike_DMS/htmls/compare_high_medium_ace2_escape.html and https://dms-vep.github.io/SARS-CoV-2_XBB.1.5_spike_DMS/htmls/compare_spike_rbd_escape.html for interactive versions of these scatter plots that also show line plots of per-site escape values for each serum.

**Extended Data Fig. 7:**
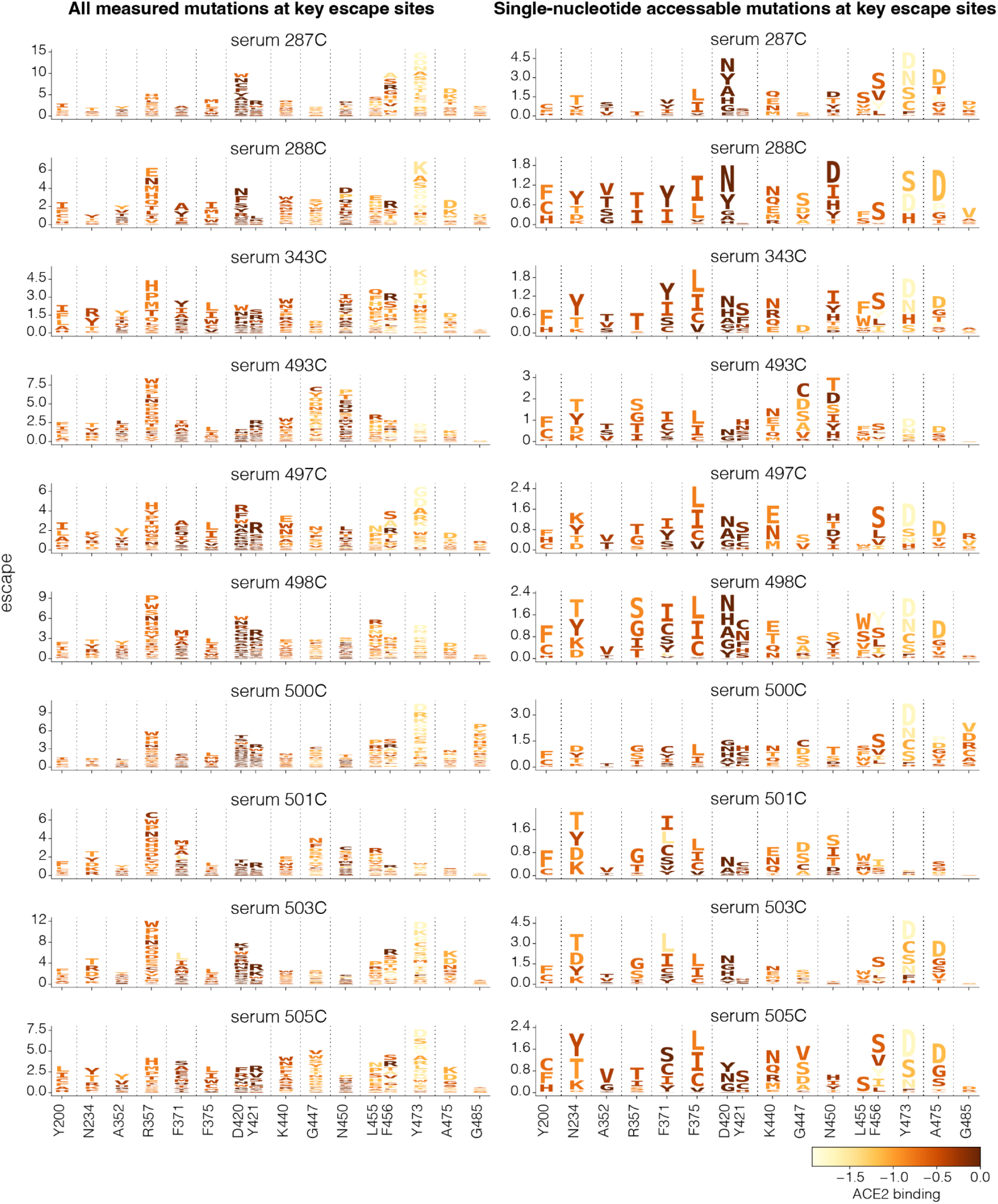
Escape at key sites for each serum. Logoplots showing XBB.1.5 spike escape at 16 highest escape sites for each of the 10 sera measured. Letter heights indicate the escape caused by mutation to that amino acid. Letters are colored light yellow to dark brown depending on mutation effect on ACE2 binding. Left: all mutations measured. Right: mutations accessible with a single-nucleotide substitution.

**Extended Data Fig. 8:**
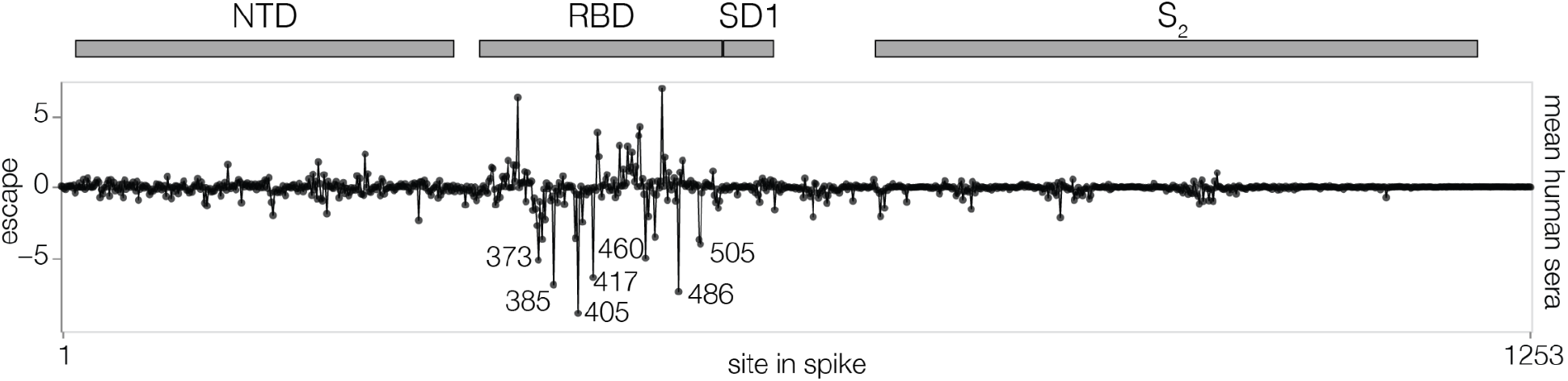
Mutations in XBB.1.5 spike that increase serum neutralization. Escape at each site in the XBB.1.5 spike averaged across the 10 sera from individuals with prior XBB* infections, showing negative as well as positive values (**Fig. 4** only shows positive values). Sites with negative escape in this plot are ones where many mutations make spike more sensitive to neutralization. Interactive plots with site and mutation-level escape are at https://dms-vep.github.io/SARS-CoV-2_XBB.1.5_spike_DMS/htmls/summary_overlaid.html (set ‘floor escape at zero’ at the bottom of the chart to false to show negative escape).

**Extended Data Fig. 9:**
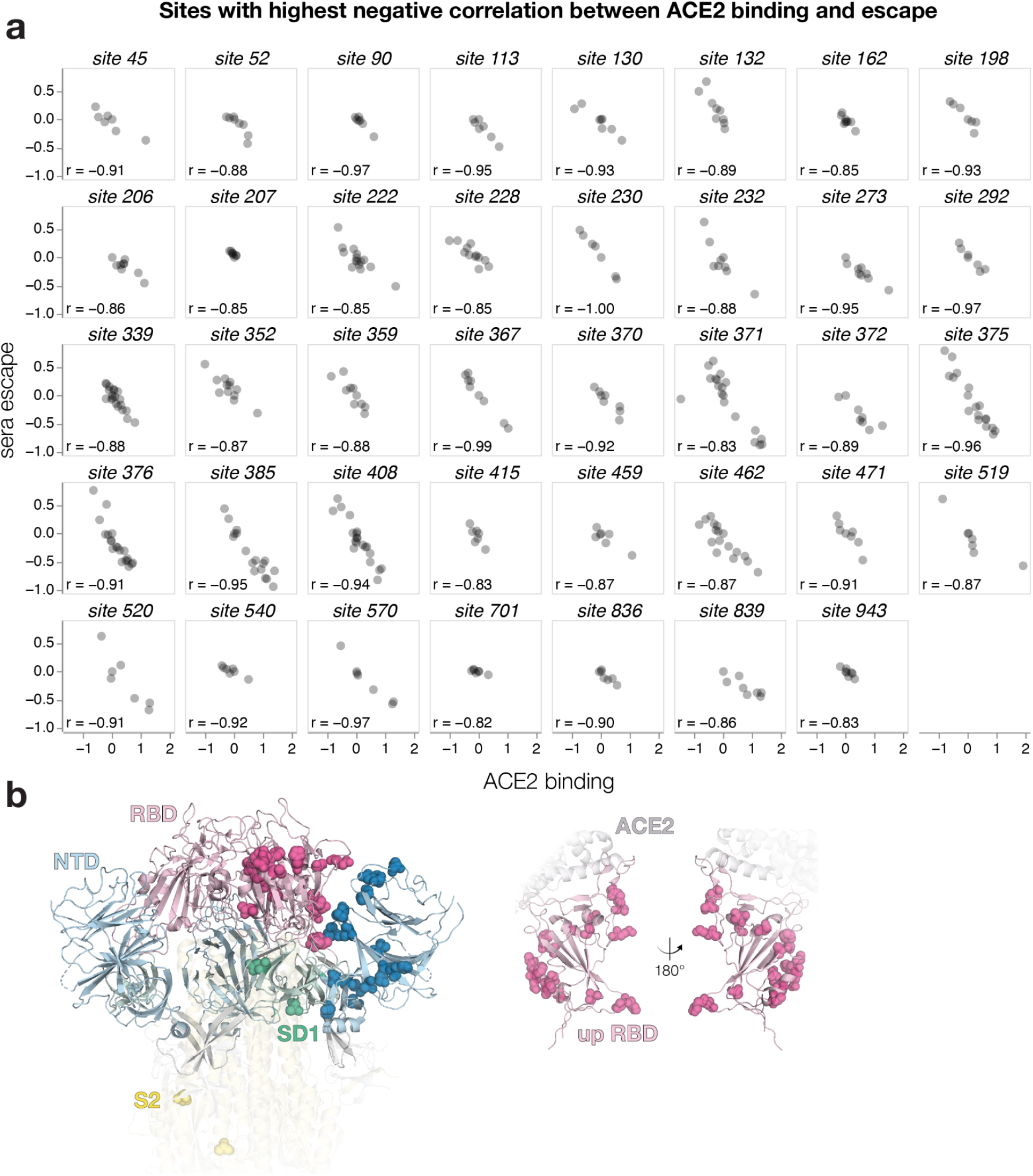
Sites with highest inverse correlation between ACE2 binding and serum escape. **a,** Correlation between ACE2 binding and serum escape for sites in XBB.1.5 spike. Only sites with at least 7 mutations measured and Pearson r < 0.82 are shown. **b,** Most sites with strongly negative correlations between mutational effects on ACE2 binding and escape are at positions that could plausibly impact the RBD conformation in the context of the full spike, since they tend to be at the interface of the RBD and other spike domains. Left: all sites from **a** shown on spike structure as spheres. RBD is colored in light pink, NTD light blue, SD1 green and the S_2_ subunit in yellow. Spheres are shown on only one chain for each domain for clarity (PDB ID: 8IOU). Right: RBD sites from **a** shown on RBD in up position engaged with ACE2. RBD is colored in light pink and ACE2 is gray.

**Extended Data Fig. 10:**
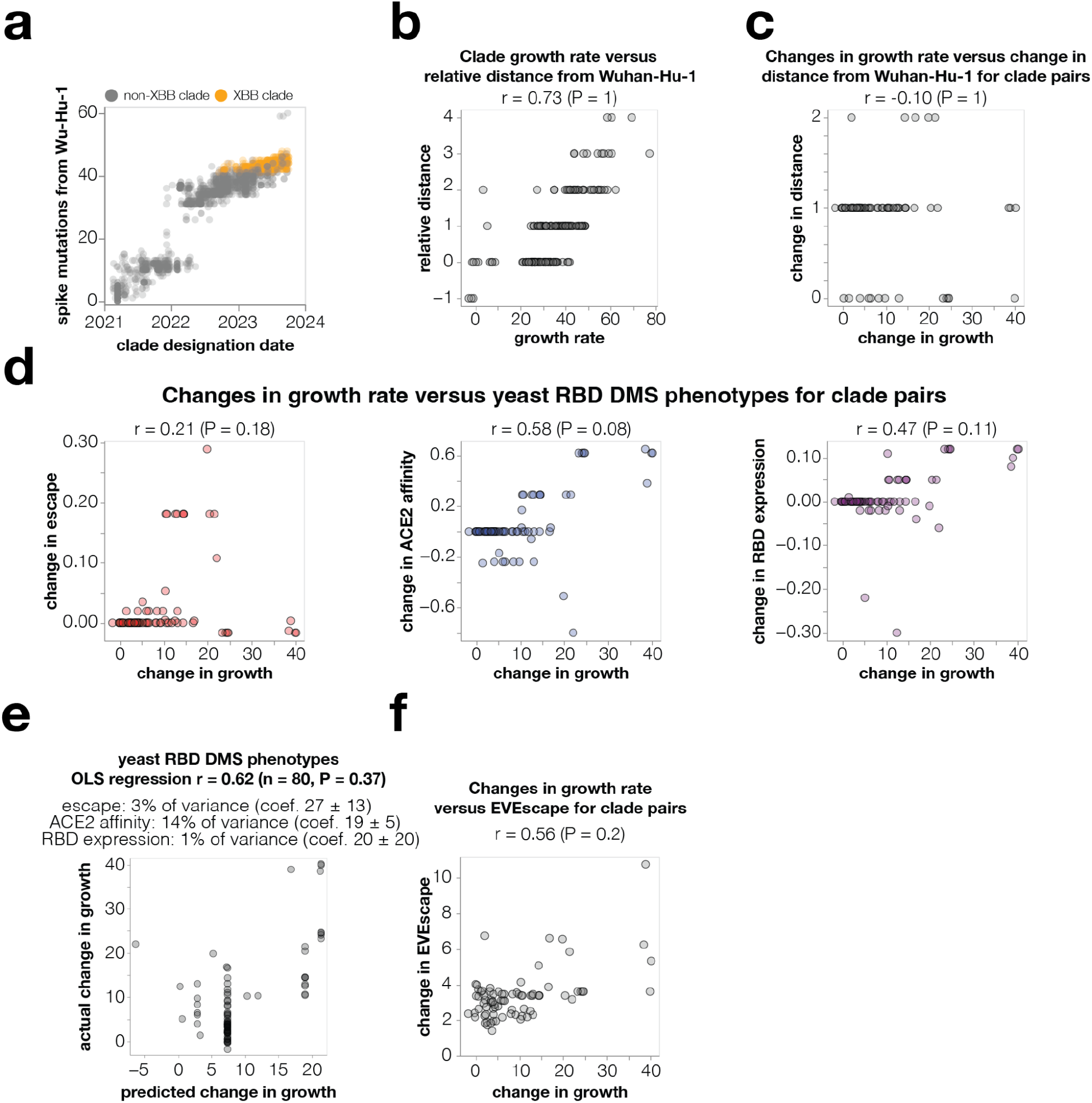
Correlations of clade growth and changes in clade growth with various other properties of spike. **a,** Number of spike amino-acid mutations relative to the early Wuhan-Hu-1 virus in all SARS-CoV-2 Pango clades versus the clade designation dates. XBB-descended clades are in orange. **b,** Because newer clades tend to have both more mutations and better growth, clade growth rate is trivially correlated with a clade’s relative distance (number of spike mutations) from Wuhan-Hu-1. However, this correlation is not informative as it is already known that new clades tend to have more mutations. **c,** If we instead correlate the change in growth rate between parent-descendant clade pairs (**Fig. 6b**) with the change in spike mutational distance to Wuhan-Hu-1 there is no correlation, since this approach removes the co-variation with total mutation count. Therefore, simple mutation counting is not informative for predicting changes in clade growth. **d,** Correlations of changes in clade growth with changes in site-level antibody escape, ACE2 affinity, and RBD expression measured for RBD mutations in yeast-display deep mutational scanning. **e,** Ordinary least-squares regression of changes in the RBD deep mutational scanning phenotypes versus changes in clade growth. **f,** Correlation of changes in the EVEscape score with changes in clade growth. Panels b-f are labeled with the Pearson correlation (r) and a P-value determined by computing how many randomizations of the mutational data yield correlations as large as the actual one.

**Extended Data Fig. 11:**
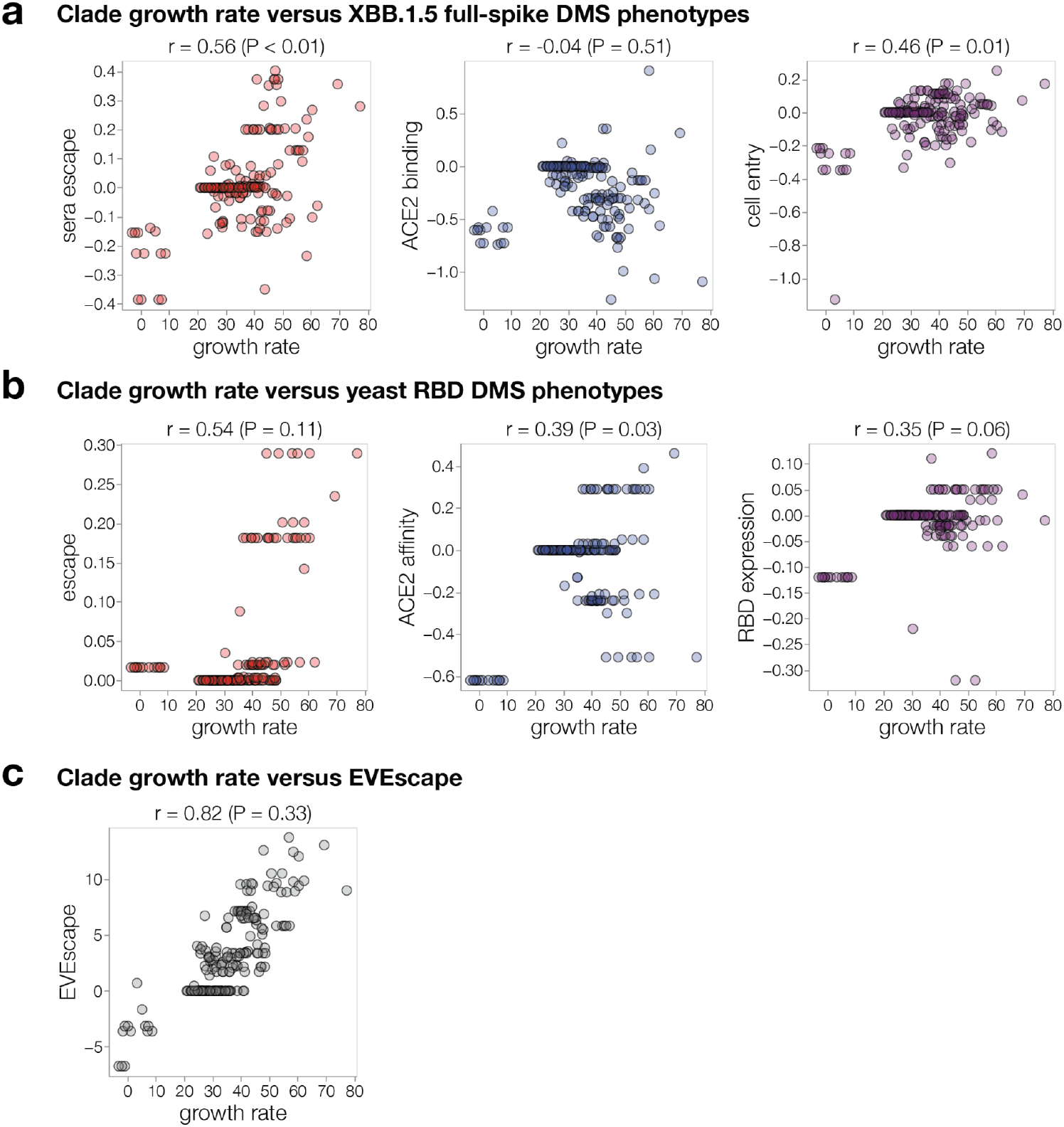
Correlations in absolute clade growth with absolute clade phenotypes. Correlations for **a,** the phenotypes measured by the full spike deep mutational scanning in the current paper; **b,** the phenotypes measured in yeast display RBD deep mutational scanning; **c,** predicted by the EVEscape method. These plots differ from **Fig. 6c** and **Extended Data Fig. 10d,f** in that they show the correlations in absolute clade growth with the absolute clade phenotypes, rather than comparing the changes in both for each parent-descendant clade pair. Absolute clade phenotypes are computed as the sum of mutation effects. The P-values above the plots are the fraction of times the correlation is greater than that for the actual data after randomizing the phenotypic effects among mutations. Note that the correlations are no longer at all reflective of the P-values for the reasons noted in the main text and **Extended Data Fig. 10c**—phylogenetic correlations, and the fact that new clades have both more mutations and higher growth so that any “phenotype” that amounts to counting mutations gives a correlation in these plots. For this reason, comparing changes in clade growth to changes in spike phenotypes as done in **Fig. 6c** and **Extended Data Fig. 10d,f** is the correct approach to test whether a method can actually predict which new clades will be successful.

**Supplementary Table 1:** Information on sera used in this study. Sera selected for this study was from individuals who either had a confirmed XBB* infection (marked by * in the table above) or had the last recorded infection during the period when XBB or its descendant lineages were the most common circulating variants in Washington state. In February 2023 70% of sequenced cased were confirmed XBB or its descendant lineages and between March and May this number grew from 88% to 97% according to the samples sequenced at University of Washington Virology labs^73^. NA indicates information not available for that individual.

| Serum sample | Sex | Race | Age | Number of infections | Number of vaccine and booster doses | 1st infection symptom date | 2nd infection symptom date | 3rd infection symptom date |
| --- | --- | --- | --- | --- | --- | --- | --- | --- |
| 287C | Female | White | 57 | 3 | 5 | Sep-2021 | May-2022 | Mar-2023 |
| 288C | Male | White | 56 | 3 | 5 | Aug-2021 | May-2022 | Mar-2023 |
| 343C | Female | Asian | 19 | 2 | 4 | Jan-2022 | Apr-2023 |  |
| 493C* | Male | White | 28 | 1 | 2 | Jan-2023 |  |  |
| 497C | Female | White | 48 | 2 | 4 | Apr-2022 | Mar-2023 |  |
| 498C* | NA | NA | NA | 1 | 5 | Dec-2022 |  |  |
| 500C | Male | Asian | 63 | 1 | 5 | Mar-2023 |  |  |
| 501C | Male | White | 53 | 1 | 5 | Mar-2023 |  |  |
| 503C | Male | White | 51 | 1 | 6 | May-2023 |  |  |
| 505C* | Female | Middle Eastern | 41 | 1 | 3 | Feb-2023 |  |  |

\* sequencing confirmed infection. 493C and 505C had XBB.1.5 infection and 498C had XBB infection.

